# Pyruvate-formate lyase–derived formate regulates the response of *Salmonella* Typhimurium to meropenem and ciprofloxacin

**DOI:** 10.1101/2025.05.22.655487

**Authors:** Debapriya Mukherjee, Sakshi Vasant Wagle, Pallab Ghosh, Abhishek Biswas, Dipshikha Chakravortty

## Abstract

*Salmonella enterica* serovar Typhimurium (STM) employs various strategies to endure antibiotic stress, but the contribution of central metabolites to resistance remains insufficiently characterized. In this work, we identify intracellular formate, produced by pyruvate-formate lyase (PflB), as a key determinant of STM susceptibility to meropenem and ciprofloxacin. Deletion of *pflB* disrupts cellular pH homeostasis, impairing efflux pump function, increasing reactive oxygen species, and causing membrane depolarization, collectively heightening sensitivity to antibiotics. Supplementing extracellular formate reverses these phenotypes. Mechanistically, pH imbalance activates the RpoE–*csrB* regulatory pathway, leading to reduced expression of efflux pump genes *acrB* and *tolC*, while extracellular formate engages the BarA/SirA two-component system to further influence the CsrA/*csrB* network. Disrupting the formate pool interferes with efflux-dependent resistance, revealing a previously unrecognized signalling function for intracellular metabolites. Overall, our results position formate as a metabolic driver of antibiotic outcomes in STM and highlight it as a potential target for mitigating resistance.

## Introduction

The global escalation of antimicrobial resistance (AMR) has been driven largely by decades of antibiotic overuse and misuse (1, 2). Among the resistant pathogens of greatest concern, multidrug-resistant (MDR) *Salmonella* has become a significant public health threat, contributing heavily to global illness and death. In 2019, AMR-associated *Salmonella* infections were estimated to cause approximately 150,000 deaths worldwide, with 23,700 of these attributed specifically to MDR strains (3). Within this landscape, *Salmonella enterica* serovar Typhimurium (STM) remains one of the most prevalent MDR serotypes, compromising the efficacy of frontline antibiotics and posing major challenges to healthcare systems (4–7).

To survive antibiotic exposure, *Salmonella* relies on a broad array of resistance mechanisms, including enzymatic degradation of antimicrobial compounds, efflux pump–mediated drug export, structural modification of antibiotics, and alteration or protection of cellular targets (8, 9). Although these classical pathways are well characterized, the influence of bacterial metabolism on shaping AMR in *Salmonella* is still poorly defined.

Our previous work identified pyruvate-formate lyase (PflB), encoded by *pflB*, as an important contributor to STM pathogenesis. PflB regulates intracellular formate levels, a short-chain fatty acid essential for maintaining intracellular pH (ipH) homeostasis. Deletion of *pflB* disturbs ipH balance, resulting in membrane stress, activation of the extra cytoplasmic sigma factor RpoE, increased expression of the small RNA *csrB*, and repression of flagellar gene expression (10). We also observed that *pflB* loss increases STM sensitivity to meropenem and ciprofloxacin, pointing to a previously unrecognized relationship between formate metabolism and antibiotic resistance.

In the present study, we systematically evaluated how intracellular formate pools and extracellular formate supplementation influence antibiotic susceptibility in STM. We show that the absence of *pflB* leads to disrupted internal pH, causing membrane damage that activates RpoE and alters expression of the efflux pump gene *acrB*. Additionally, extracellular formate modulates antibiotic resistance through the BarA/SirA two-component system and the downstream CsrA/*csrB* regulatory network, collectively contributing to reduced *acrB* expression. Together, these results demonstrate that intracellular formate directly governs STM antibiotic susceptibility, while extracellular formate exerts distinct signalling effects. Overall, our findings establish formate as a key metabolic determinant of AMR and highlight formate homeostasis as a promising target for combating MDR *Salmonella*.

## Materials and Methods

### Bacterial strains and culture conditions

The wild-type *Salmonella enterica* subsp. *enterica* serovar Typhimurium strain 14028S used in this study was kindly provided by Professor Michael Hensel (Max von Pettenkofer Institute for Hygiene and Medical Microbiology, Germany). All bacterial strains were preserved as glycerol stocks at −80 °C and revived on Luria-Bertani (LB) agar plates (HiMedia) supplemented with the appropriate antibiotics when necessary (50 µg/ml kanamycin, 50 µg/ml ampicillin, or 25 µg/ml chloramphenicol). Single colonies were inoculated into LB broth supplemented with the appropriate antibiotics and incubated overnight at 37 °C with shaking at 170 rpm. The overnight cultures were then diluted 1:100 in fresh LB medium, with or without 40 mM sodium formate (Sigma), and grown under the same conditions for 6 hours to obtain bacteria in the logarithmic growth phase. The optical density at 600 nm (OD₆₀₀) was subsequently adjusted to 0.1 in Mueller-Hinton Broth (MHB), corresponding to approximately 10⁶ colony-forming units (CFU), for use in downstream experiments. A detailed list of bacterial strains and plasmids used in this study is provided in Table 1.

**Table 1.**

| Bacterial strains/plasmids | Characteristics | Source |
| --- | --- | --- |
| *Salmonella enterica* subspecies<br>*enterica* serovar Typhimurium<br>strain 14028S | Wild type | Gift from Prof. M. Hensel |
| STM $\Delta pflB$ | Kan<sup>R</sup> | Laboratory stock |
| STM $\Delta pflB$/pQE60: *pflB* | Kan<sup>R</sup>, Amp<sup>R</sup> | Laboratory stock |
| STM $\DeltafliC$ | Kan<sup>R</sup> | Laboratory stock |
| STM $\Delta rpoE$ | Chl<sup>R</sup> | Laboratory stock |
| STM $\Delta pflB\Delta rpoE$ | Kan<sup>R</sup>, Chl<sup>R</sup> | Laboratory stock |
| STM $\Delta sirA$ | Chl<sup>R</sup> | Laboratory stock |
| STM $\Delta pflB\Delta sirA$ | Kan<sup>R</sup>, Chl<sup>R</sup> | Laboratory stock |
| pQE60 | Low copy number<br>plasmid, Amp<sup>R</sup> | Laboratory stock |
| Cip-R C1 | Ciprofloxacin resistant<br>mutant | Laboratory stock |
| Cip-R C2 | Ciprofloxacin resistant<br>mutant | Laboratory stock |
| Cip-R C3 | Ciprofloxacin resistant<br>mutant | Laboratory stock |
| Mer-R C1 | Meropenem resistant<br>mutant | Laboratory stock |
| Mer-R C2 | Meropenem resistant<br>mutant | Laboratory stock |
| Mer-R C3 | Meropenem resistant<br>mutant | Laboratory stock |
| pQE60: *pflB* | Ampicillin resistant | Laboratory stock |

### Determination of MIC of meropenem, ciprofloxacin, ceftazidime

Logarithmic phase bacterial cultures, adjusted to an OD₆₀₀ of 0.1, were inoculated into 96-well microtiter plates containing Mueller-Hinton Broth (MHB) supplemented with serial dilutions of antibiotics. The concentrations used were as follows: meropenem (8 to 0.0156 mg/L) (Sigma), ciprofloxacin (0.5 to 0.0009 mg/L) (Sigma), and ceftazidime (128 to 0.125 mg/L). Plates were incubated at 37 °C with shaking at 170 rpm for 14-16 hours. Bacterial growth was assessed by measuring OD₆₀₀ using a Tecan microplate reader.

### Estimation of bacterial viability using resazurin assay

Following antibiotic exposure and incubation for 14-16 hours, 20 µL of resazurin solution (prepared from a 0.2 mg/mL stock) was added to each well and incubated for 2 hours in the dark. Thereafter, fluorescence intensity was measured using a Tecan microplate reader with excitation at 540 nm and emission at 590 nm. Cell viability was assessed by monitoring the color change of resazurin. A stronger purple coloration indicated higher cell viability.

### Flow cytometry mediated quantification of live/dead population, intracellular reactive oxygen species (ROS), membrane depolarization, and ipH

After 14–16 hours of antibiotic treatment, bacterial cells were stained with a panel of fluorescent dyes: Propidium Iodide (PI, 1 µg/mL) for live/dead discrimination, dichlorodihydrofluorescein diacetate (DCF_2_DA, 10 µM) to assess intracellular reactive oxygen species (ROS), Bis-(1,3-Dibutylbarbituric Acid) Trimethine Oxonol (DiBAC_4_, 1 µg/mL) for measuring membrane depolarization, and 2′,7′-Bis-(2-Carboxyethyl)-5-(and-6)-Carboxyfluorescein Acetoxymethyl Ester (BCECF-AM, 20 µM) for intracellular pH (ipH) comparison. Stained samples were incubated in the dark for 15 minutes and subsequently analyzed using the BD FACSVerse flow cytometer (BD Biosciences, USA). Data including dot plots, density plots, median fluorescence intensity (MFI), and the percentage of positively stained cells were acquired and processed using BD FACSVerse software.

### Determination of intracellular formate and pyruvate concentration

Following exposure of logarithmic-phase cultures of STM WT and passaged strains to sub-lethal antibiotic concentrations and a 16-hour incubation, the cultures were centrifuged at 6,000 rpm for 10 minutes. The resulting pellet was washed twice with 1X PBS to remove residual media and then resuspended in PBS. A 100 µl portion of this suspension was plated on LB agar to determine the CFU count. The remaining suspension was lysed by sonication, and cellular debris were cleared by centrifugation at 12,000 g for 10 minutes. Formate and pyruvate levels in the supernatant were quantified using the Abcam formate estimation kit (ab111748) and Elabscience pyruvate estimation kit (E-BC-K130-S) according to the manufacturer’s instructions, and the concentrations were interpolated from a standard curve. The concentrations were normalized with the CFU obtained from plating.

### RNA isolation and RT-qPCR

Logarithmic-phase bacterial cultures were harvested by centrifugation at 6,000 rpm for 10 minutes at 4 °C, and the resulting cell pellets were resuspended in TRIzol reagent. Samples were stored at −80 °C until further processing. Total RNA was extracted using the chloroform–isopropanol method, and RNA pellets were washed with 70% ethanol and resuspended in 30 µL of DEPC-treated Milli-Q water. RNA concentration was measured using a NanoDrop spectrophotometer (Thermo Fisher Scientific), and RNA integrity was verified by electrophoresis on a 2% agarose gel. For removal of genomic DNA, 2 µg of total RNA was treated with TURBO DNase (Thermo Fisher Scientific) at 37 °C for 30 minutes, followed by inactivation with 5 mM Na₂EDTA at 65 °C for 10 minutes. First-strand cDNA synthesis was performed using the PrimeScript RT reagent kit (Takara, Cat. No. RR037A) according to the manufacturer’s protocol. Quantitative real-time PCR (qRT-PCR) was carried out using the TB Green RT-qPCR kit (Takara) on a QuantStudio 5 Real-Time PCR System. Gene expression was analysed using intragenic primers and normalized to *16S rRNA* levels. Primer sequences are provided in Table 2.

**Table 2.**
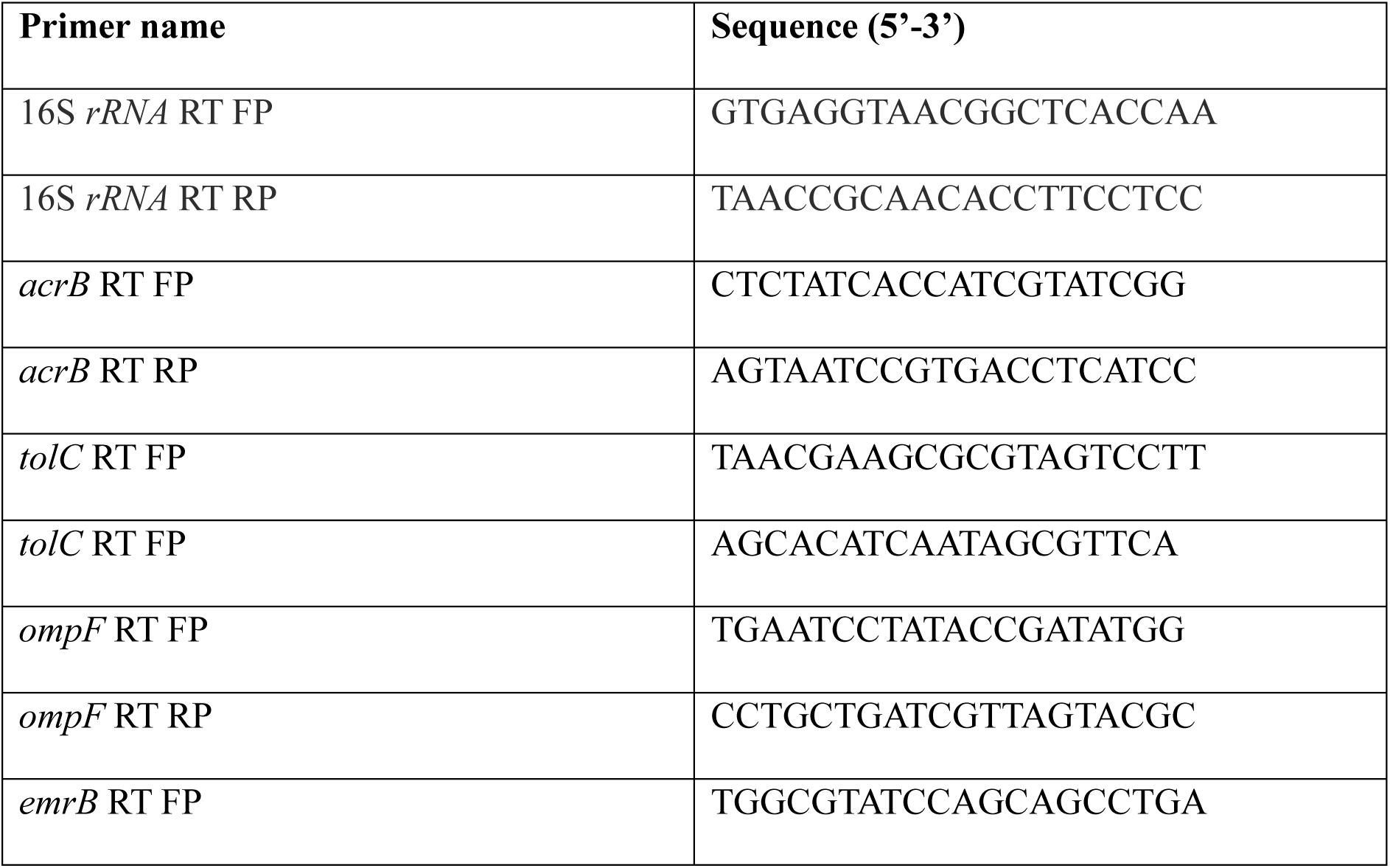
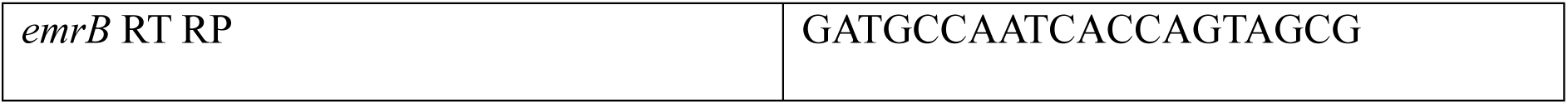

### Efflux pump activity by Nile Red Assay

Efflux pump activity was quantified following the method outlined by Lyu *et al*(11). In brief, bacterial cultures in the logarithmic growth phase were adjusted to an optical density of 0.1 at 600 nm (OD₆₀₀). A volume of 100 µl from each normalized culture was dispensed into wells of a 96-well microplate and incubated with Nile Red dye at a final concentration of 48 µg/ml. The plate was incubated in the dark under shaking conditions for 3 hours. Fluorescence measurements were then obtained using a TECAN microplate reader, with excitation and emission wavelengths set at 549 nm and 628 nm, respectively. Fluorescence intensity values were normalized against absorbance readings at 600 nm.

### Atomic Force Microscopy

After 14–16 hours of antibiotic exposure, bacterial cells were drop-cast onto sterile glass coverslips. Once air-dried, the cells were fixed by treating them with 3.5% paraformaldehyde for 10 minutes. Residual salts were removed by washing the samples three times with double-autoclaved MilliQ water. The coverslips were then dried in a vacuum desiccator prior to imaging with the NX10 Atomic Force Microscope (AFM). Image processing and measurements of cell length and height were performed using XEI software.

### Animal experiments

All animal experiments were conducted with prior approval from the Institutional Animal Ethics Clearance Committee (IAEC) at the Indian Institute of Science, Bangalore. The institute is registered under CPCSEA with registration number 48/1999/CPCSEA. All procedures strictly adhered to the guidelines set by the Committee for the Purpose of Control and Supervision of Experiments on Animals (CPCSEA). The ethical approval number for this study is CAF/Ethics/853/2021.

A modified protocol based on the method described by Roy Chowdhury *et al.* was employed to evaluate the organ burden following infection and antibiotic treatment with the wild-type (WT) and mutant bacterial strains(12). Briefly, 10⁶ cells from overnight cultures were orally gavaged into 6–8-week-old C57BL/6 mice on day 0 (0 dpi). Mice received 1 mg/kg of the respective antibiotics on days 2 and 4 post-infection. On day 5 post-infection, the animals were ethically euthanized, and the intestine, liver, and spleen were collected. These organs were homogenized, and serial dilutions of the homogenates were plated on *Salmonella-Shigella* (SS) agar. Bacterial load was quantified and expressed as log₁₀ CFU per gram of tissue.

To compare mortality rates between STM WT and STM Δ*pflB* under antibiotic exposure, groups of mice (n=5) were orally administered 10⁶ cells from stationary phase cultures of each strain. Starting from day 2 post-infection, 1 mg/kg of the respective antibiotics was administered every alternate day till the mice survived. Survival was monitored daily and represented as percent survival. A 125 mM solution of sodium formate was supplemented in the drinking water provided to the animals in both experimental setups.

### Estimation of Formate in mouse serum by Gas Chromatography-Mass Spectrometry

To quantify formate levels in mouse serum following supplementation with 125 mM sodium formate in drinking water, a protocol adapted from Hughes *et al* was employed(13). Male C57BL/6 mice (6–8 weeks old) were provided with 125 mM sodium formate in their drinking water. At five days post-infection, blood was collected via retro-orbital puncture. After clotting, serum was separated by centrifugation at 10,000 × g for 20 minutes and subsequently used for formate quantification.

Sodium formate (Sigma) served as an external standard. Both standards and biological samples were derivatized prior to mass spectrometry analysis. To protonate the formate, 2 M hydrochloric acid was added to each sample in a 1:1 ratio. Liquid-liquid extraction was performed to isolate formic acid into ethyl acetate (Sigma-Aldrich). Residual water was removed by passing the ethyl acetate extract through an anhydrous sodium sulfate column (Sigma-Aldrich). The extract was then incubated at 80 °C for 1 hour with the derivatizing reagent N,O-Bis(trimethylsilyl)trifluoroacetamide (Sigma-Aldrich) containing 1% chlorotrimethylsilane (Fluka). Derivatized samples were transferred to autosampler vials for analysis using Gas Chromatography-Mass Spectrometry (GC-MS; Agilent Technologies, 7890A GC, 5975C MS).

Injection parameters were as follows: injection temperature of 200 °C, injection split ratio of 1:40, and injection volume of 1 µL. The oven temperature program began at 40 °C (held for 4 minutes), increased to 120 °C at a rate of 5 °C/min, and then to 200 °C at 25 °C/min, with a final hold of 3 minutes. Helium was used as the carrier gas at a constant flow rate of 1.2 mL/min. Chromatographic separation was performed using a 30 m × 0.25 mm × 0.40 µm DB-624 column (Agilent). The interface temperature was maintained at 220 °C, and the electron impact ion source operated at 230 °C with an ionization energy of 70 eV and 1235 EM volts.

For qualitative analysis, selected ion monitoring in single quadrupole mode was employed with a scan at m/z 103. The retention time for formate and its deuterated analog was approximately 5.9 minutes. The target and reference (qualifier) ions for formate were m/z = 103 and m/z = 75, respectively. Sample concentrations were determined by interpolating from a standard calibration curve generated using known concentrations.

### Determination of the fraction of antibiotic persister

The determination of antibiotic persister was performed by modifying a protocol described by Drescher *et al*(14). Logarithmic phase cultures of STM WT and STM Δ*pflB* strains were treated with 15X and 25X the minimum inhibitory concentration (MIC) of meropenem (2 and ciprofloxacin. To determine the persister population, aliquots were collected at specified timepoints post-treatment and plated for colony-forming unit (CFU) enumeration. Similarly, 24-hour-old biofilm cultures were exposed to the same antibiotic concentrations, and persister cells were quantified via plating. The resulting CFU counts were normalized against the pre-inoculum values to calculate fold viability relative to the 0-hour timepoint. Data were expressed as log₁₀(fold decrease in CFU relative to 0 h).

### Statistical Analysis

Each experiment was independently repeated between 1 and 12 times, as specified in the figure legends. Statistical analyses were conducted using an unpaired Student’s t-test, or two-way ANOVA, as indicated in the respective legends. Data from mouse experiments were analysed using the Mann–Whitney U test. A p-value of less than 0.05 was considered statistically significant. Results are presented as Mean ± SD or Mean ± SEM, as noted in the figure legends. All data analysis and graphical representations were performed using GraphPad Prism version 8.4.2.

## Results

### *pflB* was transcriptionally upregulated in ciprofloxacin resistant mutants, indicating a possible involvement of intracellular formate in regulating *Salmonella* AMR

In our earlier work, we demonstrated that *pflB* plays a central role in maintaining ipH in *Salmonella*, which in turn governs the adhesion–invasion switch during its pathogenic lifestyle (10). Other studies have also highlighted the importance of ipH in shaping bacterial antimicrobial responses (15, 16). These observations prompted us to investigate whether intracellular formate levels might also contribute to the regulation of AMR in *Salmonella*.

To investigate formate-associated responses, we analysed *pflB* expression in ciprofloxacin-resistant mutants (Cip-R C1, C2, C3) generated by passaging STM for 160 generations under sublethal ciprofloxacin. All Cip-R strains showed increased *pflB* transcription relative to WT **(Figure 1A)**. In Cip-R C1, sublethal ciprofloxacin further improved *pflB* expression **(Figure 1B)**, which coincided with elevated intracellular formate levels under drug exposure **(Figure 1C)**.

**Figure 1:**
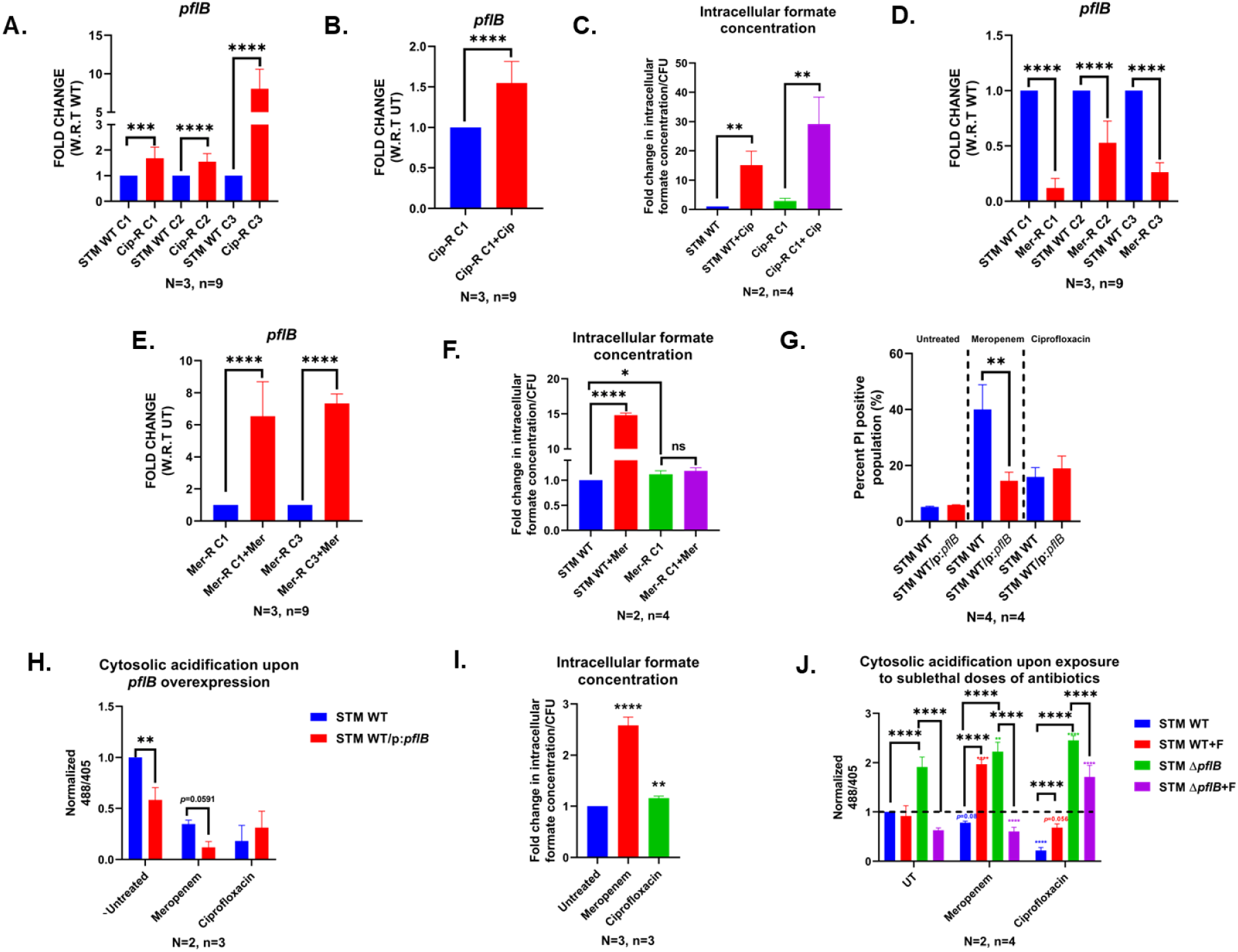
Higher *pflB* transcription in the presence of antibiotics was frequently observed as a characteristic of ciprofloxacin and meropenem mutants, and loss of *pflB* was linked to impaired acid homeostasis when exposed to sublethal concentrations of meropenem and ciprofloxacin. (A) Transcriptional upregulation of *pflB* in Cip-R 1, 2, 3 ciprofloxacin resistant STM strains. Data representative of N=3, n=9 and is presented as mean+/- SD. (B) Transcriptional upregulation of *pflB* in Cip-R C1 upon exposure to sub-lethal doses of ciprofloxacin. Data representative of N=3, n=9 and is presented as mean+/- SD. (C) Quantification of intracellular formate levels in STM WT and Cip-R C1 strains upon exposure to sub-lethal dose of ciprofloxacin. Data representative of N=2, n=4 and is presented as mean+/- SD. (D) Transcriptional levels of *pflB* in Mer-R C1, 2, 3 meropenem resistant STM strains. Data representative of N=3, n=9 and is presented as mean+/- SD. (E) Transcriptional upregulation of *pflB* in Mer-R C1, 3 upon exposure to sub-lethal doses of ciprofloxacin. Data representative of N=3, n=9 and is presented as mean+/- SD. (F) Quantification of intracellular formate levels in STM WT and Mer-R C1 strains upon exposure to sub-lethal dose of ciprofloxacin. Data representative of N=2, n=4 and is presented as mean+/- SD. (G) Bar graphs showing PI-positive population in STM WT and STM WT/pQE60:*pflB* upon exposure to meropenem and ciprofloxacin. Data representative of N=2, n=4 and is presented as mean+/- SD. (H) Normalized 488/405 ratio of STM WT and STM WT/pQE60:*pflB* in presence of sub-lethal doses of meropenem and ciprofloxacin. Data representative of N=2, n=4 and is presented as mean+/- SD. (I) Quantification of intracellular formate levels in STM WT upon exposure to sub-lethal dose of ciprofloxacin and meropenem. Data representative of N=3, n=3 and is presented as mean+/-SD. (J) Normalized 488/405 ratio of STM WT and STM Δ*pflB* in presence of sub-lethal doses of meropenem and ciprofloxacin. Data representative of N=2, n=4 and is presented as mean+/-SD. (Unpaired two-tailed Student’s t-test for column graphs, Two-way ANOVA for grouped data, Mann-Whitney U-test for animal experiment data (*** p < 0.0001, *** p < 0.001, ** p<0.01, * p<0.05))

In contrast, meropenem-resistant strains (Mer-R C1, C2, C3), obtained via sublethal meropenem passaging, displayed reduced *pflB* transcription **(Figure 1D)**. However, Mer-R C1 and Mer-R C3 upregulated *pflB* upon sublethal meropenem exposure **(Figure 1E)**. Consistently, Mer-R C1 had a higher intracellular concentration formate than the WT strain **(Figure 1F)**. Together, these results indicate that elevated intracellular formate is a recurring feature of strains evolved under ciprofloxacin or meropenem pressure.

Since *pflB* encodes pyruvate-formate lyase, the major source of intracellular formate in *Salmonella*, we used cloned *pflB* in the plasmid pQE60 and overexpressed it using Isopropyl β-D-1-thiogalactopyranoside (IPTG) to assess the functional impact of enhanced *pflB* (17). In STM WT, *pflB* overexpression reduced the Propidium Iodide (PI)–positive population—signifying lower cell death—after meropenem treatment, with no effect observed under ciprofloxacin exposure **(Figure 1G)**. *pflB* overexpression also decreased intracellular pH in both untreated and meropenem-treated cells **(Figure 1H)**, consistent with our earlier observation that *pflB* deletion disturbs pH homeostasis.

Aligned with these observations, meropenem and ciprofloxacin exposure continued to increase intracellular formate in our laboratory WT strain **(Figure 1I)**. To probe the role of ipH in antibiotic responses, we monitored ipH changes in STM WT and STM Δ*pflB* following sublethal meropenem or ciprofloxacin treatment using the ratiometric dye BCECF-AM. STM WT exhibited a reduced 488/405 ratio under antibiotic stress, indicating cytoplasmic acidification. STM Δ*pflB*, which shows a higher basal ratio, did not acidify upon antibiotic exposure, suggesting resistance to pH drops. Formate supplementation partially rescued the Δ*pflB* phenotype by decreasing the 488/405 ratio under antibiotic stress. Although formate alone had little effect on WT ipH, antibiotic-treated WT displayed an increased 488/405 ratio with formate **(Figure 1J)**. Since PflB converts pyruvate into formate, we found that *pflB* deletion resulted in elevated intracellular pyruvate levels under untreated conditions. These higher pyruvate levels in STM Δ*pflB* persisted during meropenem and ciprofloxacin exposure. In contrast, STM WT strain showed an overall increase in pyruvate upon antibiotic treatment, suggesting that the cell may upregulate *pflB* to channel pyruvate toward formate production as a strategy to counter antibiotic stress by lowering intracellular pH **(Figure S1A)**.

Together, these findings indicate that formate is a key metabolite whose intracellular levels commonly rise during antibiotic exposure, and whose depletion disrupts pH homeostasis under antibiotic stress.

### Deletion of *pflB* enhances susceptibility of STM to meropenem and ciprofloxacin, whereas supplementation of extracellular formate also heightens susceptibility of STM to these antibiotics

To investigate the impact of deleting *pflB*, which was consistently upregulated during antibiotic exposure, we examined how this gene deletion affected the antibiotic susceptibility of STM. To assess the sensitivity of the STM Δ*pflB* strain to meropenem and ciprofloxacin, both the wild-type and mutant strains were treated with a range of antibiotic concentrations. The STM Δ*pflB* mutant showed significantly increased sensitivity to meropenem, particularly at concentrations of 1 mg/L and 0.5 mg/L **(Figure 2A)**. While the partial restoration of bacterial growth upon sodium formate supplementation was not statistically significant, it suggests that intracellular formate deficiency may contribute to the heightened sensitivity to meropenem. Interestingly, when formate was added to wild-type cells—despite their intact intracellular formate levels—a significant reduction in survivability was observed under antibiotic stress at the same concentrations. Comparable results were obtained with ciprofloxacin, where statistically significant effects were noted at 0.03125, 0.015625, 0.0078, and 0.0039 mg/L **(Figure 2B)**. These findings were further supported by resazurin assay data **(Figure 2C, D)**. The enhanced susceptibility of STM Δ*pflB* and its recovery in the presence of sodium formate was also true for ceftazidime, a third generation cephalosporin (18) **(Figure S1B)**.

**Figure 2:**
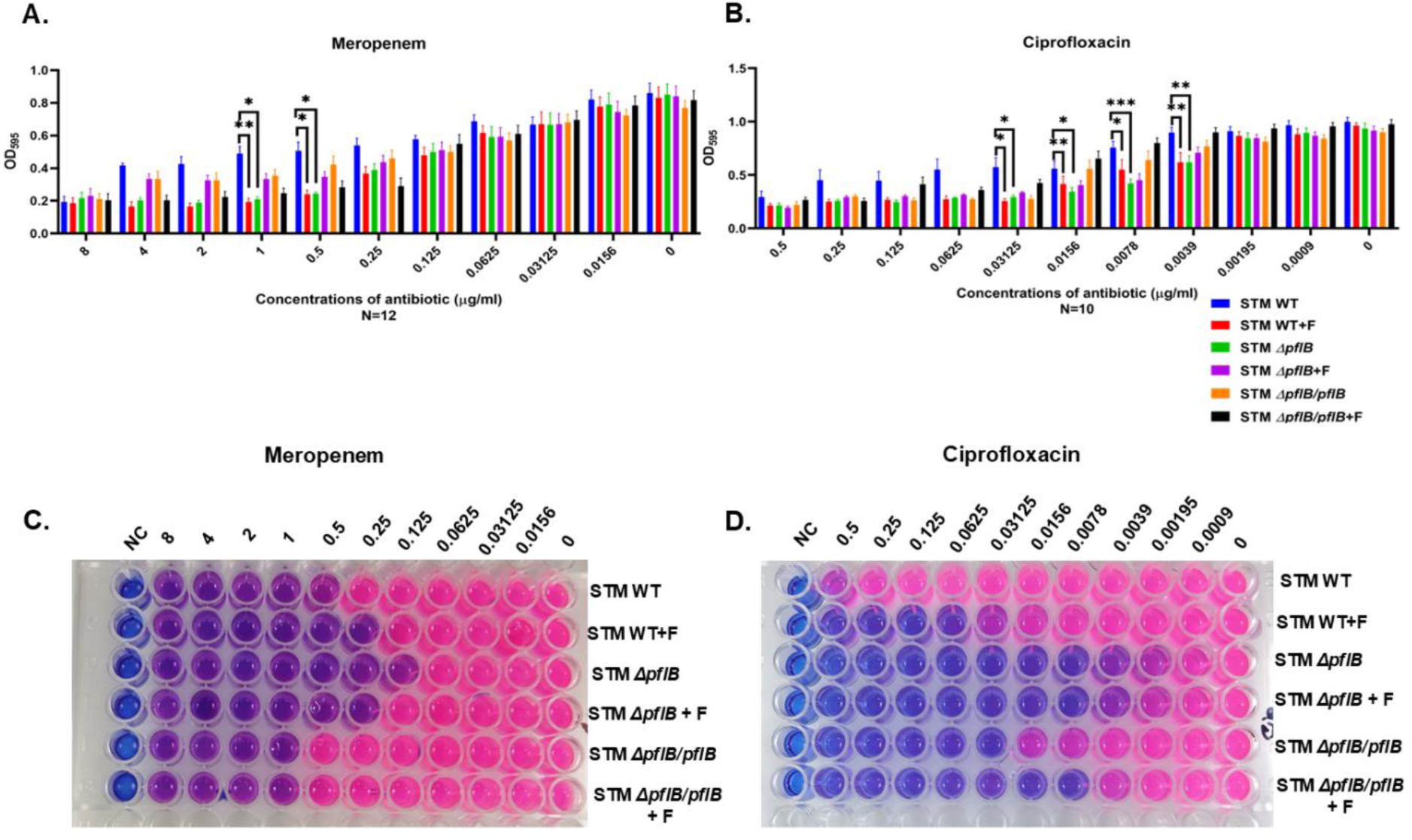
Deletion of *pflB* increases STM’s sensitivity to meropenem and ciprofloxacin, while extracellular formate supplementation similarly enhances antibiotic susceptibility. (A) Meropenem susceptibility of STM WT, STM Δ*pflB*, and STM Δ*pflB*/pQE60: *pflB*, with or without formate treatment, was assessed using the microfold dilution method. Data presented as mean+/- SEM of N=12. (B) Ciprofloxacin susceptibility of STM WT, STM Δ*pflB*, and STM Δ*pflB*/pQE60: *pflB*, with or without formate treatment, was assessed using the microfold dilution method. Data presented as mean+/- SEM of N=10. (C) Resazurin assay of STM WT, STM Δ*pflB*, and STM Δ*pflB*/pQE60: *pflB*, with or without formate treatment, in increasing concentrations of meropenem. (D) Resazurin assay of STM WT, STM Δ*pflB*, and STM Δ*pflB*/pQE60: *pflB*, with or without formate treatment, in increasing concentrations of ciprofloxacin. (Unpaired two-tailed Student’s t-test for column graphs, Two-way ANOVA for grouped data, Mann-Whitney U-test for animal experiment data (*** p < 0.0001, *** p < 0.001, ** p<0.01, * p<0.05))

To investigate how deletion of *pflB* affects STM’s antibiotic susceptibility, we treated logarithmic-phase cultures overnight with sub-lethal doses of meropenem (0.5 mg/L) and ciprofloxacin (0.015625 mg/L)and analyzed PI uptake via flow cytometry. STM Δ*pflB* showed a higher PI-positive population than WT, which was significantly reduced with sodium formate supplementation. In contrast, formate supplementation increased PI-positive cells in WT upon antibiotic exposure **(Figure 3A, B)**.

**Figure 3:**
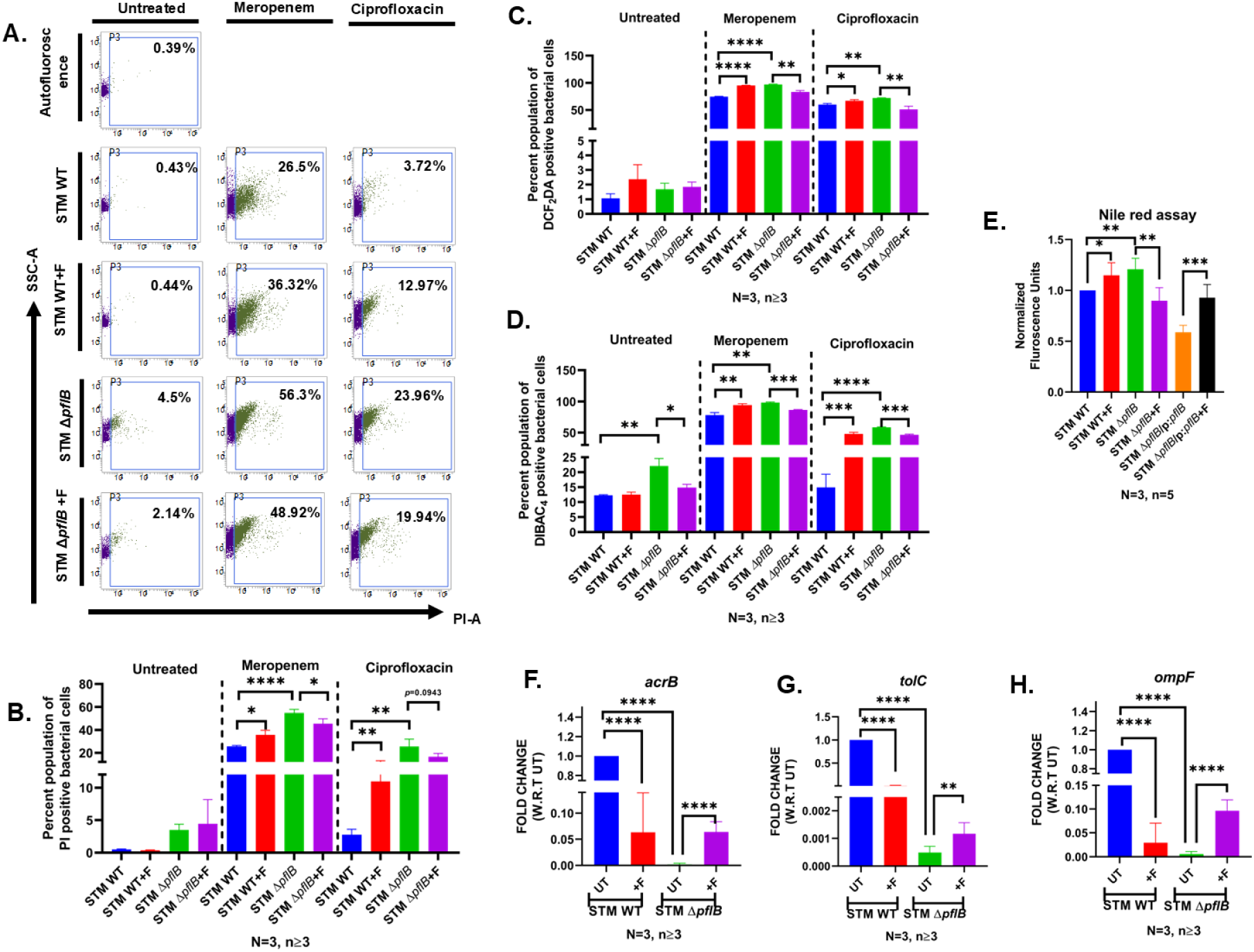
Increased antibiotic sensitivity following *pflB* deletion was linked to transcriptional downregulation of efflux pumps, a pattern also observed in formate-treated wild-type cells. (A) Dot plots (SSC-A vs PI-A) and bar graph illustrate increased bacterial killing with meropenem and ciprofloxacin following *pflB* deletion and formate supplementation in STM WT. Data representative of N=3, n≥3. (B) Bar graphs depict the PI-positive population after exposure to meropenem and ciprofloxacin following *pflB* deletion and formate supplementation in STM WT. Data represent N=3, n≥3 and are shown as mean ± SD. (C) Bar graphs display the DiBAC_4_-positive population under meropenem and ciprofloxacin treatment after *pflB* deletion and formate supplementation in STM WT. Data represent N=3, n≥3 and are shown as mean ± SD. (D) Bar graphs present the DCF_2_DA-positive population upon exposure to meropenem and ciprofloxacin following *pflB* deletion and formate supplementation in STM WT. Data represent N=3, n≥3 and are shown as mean ± SD. (E) Nile red assay quantifies efflux pump activity in STM WT, STM Δ*pflB, and* STM Δ*pflB*/pQE60:*pflB .* Data representative of N=3, n=5 and is presented as mean+/- SD. (F-H) Transcript levels of *acrB*, *tolC*, and *ompF* measured in STM WT and STM Δ*pflB*, with or without formate treatment. Data representative of N=3, n≥3 and is presented as mean+/- SD. (Unpaired two-tailed Student’s t-test for column graphs, Two-way ANOVA for grouped data, Mann-Whitney U-test for animal experiment data (*** p < 0.0001, *** p < 0.001, ** p<0.01, * p<0.05))

We next assessed whether *pflB* deletion and formate supplementation affected susceptibility to additional antibiotics. STM Δ*pflB* displayed increased sensitivity to sub-lethal doses of the fluoroquinolone levofloxacin (1 µg/ml), the carbapenem doripenem (1 µg/ml), and the β-lactam ampicillin (64 µg/ml), with partial restoration of resistance following formate supplementation (19–21). Formate addition to STM WT also heightened susceptibility to these antibiotics **(Figure S2A–C)**. These observations suggest that the increased antimicrobial sensitivity caused by *pflB* deletion or by external formate supplementation may extend beyond meropenem and ciprofloxacin. Nevertheless, to maintain focus and clarity, we concentrated on meropenem and ciprofloxacin in the subsequent analyses.

Given that bactericidal antibiotics induce reactive oxygen species (ROS) irrespective of their mechanisms, we assessed intracellular ROS using DCF_2_DA staining(22–24). Consistent with PI staining results, STM Δ*pflB* exhibited elevated DCF_2_DA-positive populations under antibiotic stress, which was mitigated by formate. Similarly, formate supplementation increased ROS levels in WT cells exposed to antibiotics **(Figure 3C, S3 A)**. These findings suggest that loss of *pflB*, and thus intracellular formate, increases STM’s susceptibility to meropenem and ciprofloxacin, while exogenous formate to wild-type cells also enhances antibiotic sensitivity.

### Deletion of *pflB* was associated with a transcriptional downregulation of antibiotic efflux pumps, which was also common for formate supplemented in wild-type cells

Our previous work demonstrated increased membrane depolarization in STM Δ*pflB* compared to WT cells (10). Previous studies have shown that outer membrane of Gram-negative pathogens forms a robust permeability barrier which can prevent antibiotics to reach their target (25). Given that antimicrobial agents like antimicrobial peptides can exert bactericidal effects via membrane depolarization we hypothesized that the enhanced antibiotic susceptibility of STM Δ*pflB* may be attributable, at least in part, to its depolarized membrane state(26–30). Notably, membrane depolarization was further exacerbated upon exposure to sub-lethal concentrations of meropenem and ciprofloxacin, while sodium formate partially alleviated this effect in STM Δ*pflB*. In contrast, formate supplementation had no impact on membrane potential in WT cells under unstressed conditions, although increased depolarization was observed upon antibiotic exposure **(Figure 3D, S3 B)**.

Efflux-mediated resistance represents a major mechanism of antibiotic failure in Gram-negative bacteria. *Salmonella* harbors multiple efflux systems, among which the TolC-dependent resistance-nodulation-division (RND) family plays a central role in expelling a broad range of clinically relevant antibiotics(31–34). As proton motive force (PMF) and pH homeostasis are critical for the functioning of these pumps, and membrane depolarization can disrupt PMF, we hypothesized that the reduced antibiotic efflux in STM Δ*pflB* may stem from impaired PMF(11). To test this, we employed a nile red efflux assay. STM Δ*pflB* retained significantly higher levels of nile red fluorescence, indicating decreased efflux activity. Formate supplementation reduced nile red retention, suggesting partial restoration of efflux capacity. Similarly, formate-treated STM WT cells exhibited increased nile red fluorescence upon antibiotic exposure, consistent with reduced efflux function **(Figure 3E)**.

To determine whether reduced efflux activity was due solely to membrane depolarization or also involved transcriptional regulation, we measured mRNA levels of *acrB* and *tolC*, two key components of the RND efflux system in *Salmonella*(35). Deletion of *pflB* resulted in transcriptional downregulation of both genes, with partial recovery upon formate supplementation. Interestingly, formate supplementation in STM WT also led to reduced *acrB* and *tolC* expression **(Figure 3F, G)**. These findings suggest that while membrane depolarization may directly impair efflux pump activity, additional transcriptional regulatory mechanisms may also contribute to reduced efflux and heightened antibiotic susceptibility.

Outer membrane proteins (Omps) influence antibiotic susceptibility in *Salmonella*, with reduced OmpF expression typically linked to resistance (36–38). Notably, *ompF* was downregulated in STM Δ*pflB* **(Figure 3H)**, even though this strain showed increased antibiotic sensitivity. This indicates that the heightened susceptibility of STM Δ*pflB* occurs independently of Omps, as decreased OmpF would normally correlate with resistance rather than greater sensitivity.

### Formate supplementation in drinking water and intrabacterial formate deficiency both enhanced bacterial clearance by antibiotics *in-vivo*

We tested whether our *in-vitro* results also applied in living animals by giving C57BL/6 mice an oral dose of 10⁶ CFU of STM WT or STM Δ*pflB*. The mice also received 125 mM sodium formate in their drinking water **(Figure 4A)**. Earlier work suggests that mice need about 40 mmol of formate per day. Since C57BL/6 mice usually drink around 30 ml of water daily, they would need about a 1 M solution to reach that amount. However, previous studies showed that 500 mM sodium formate can cause weight loss, so we selected a safer dose of 125 mM, representing about 10% of the estimated daily requirement (39). In our pilot tests, this amount was enough to significantly raise serum formate levels by 5 days post-infection **(Figure S4A)**. Our findings showed that adding formate to the drinking water together with meropenem treatment lowered bacterial loads more than meropenem alone in the intestine, liver, and spleen of C57BL/6 mice. As expected, the STM Δ*pflB* mutant showed reduced organ colonization (10); but formate supplementation did not restore its numbers. Meropenem further reduced the bacterial load, and—unlike our *in-vitro* results—formate did not rescue the decreased bacterial counts. Interestingly, combining formate with meropenem did not lower organ colonization in STM Δ*pflB* the way it did in STM WT, suggesting that STM Δ*pflB* may use extracellular formate to survive antibiotic stress (**Figure 4B**). The same trends were seen with ciprofloxacin (**Figure 4C**) and ceftazidime (**Figure 4D**) in mice infected with STM WT or STM Δ*pflB* under formate-supplemented conditions.

**Figure 4:**
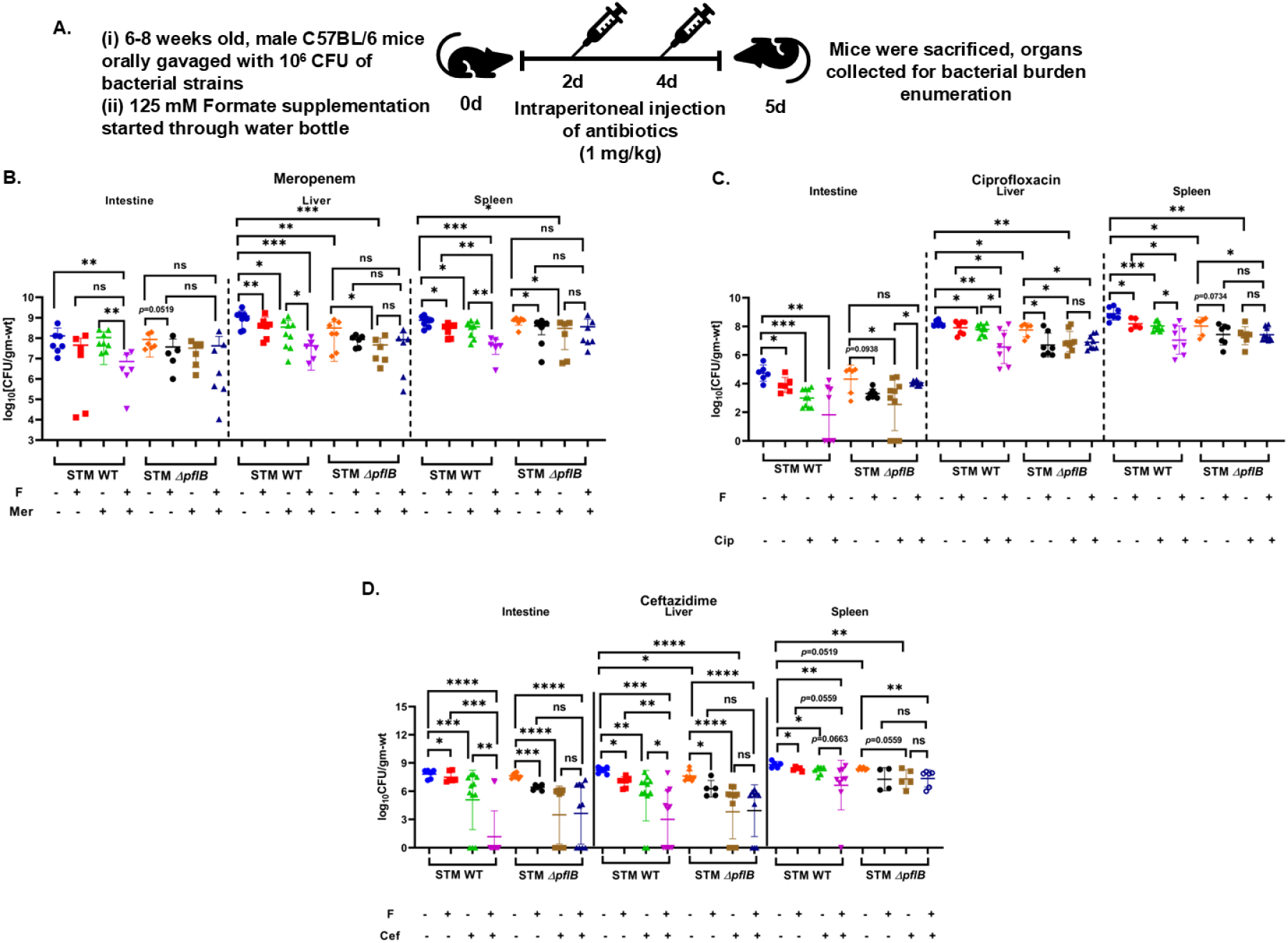
Both formate supplementation via drinking water and internal formate deficiency enhanced antibiotic-mediated bacterial clearance *in vivo*. (A) Schematic outlining the experimental design. (B-D) Role of formate supplementation in STM WT and STM Δ*pflB* clearance from the intestine, liver, and spleen following treatment with meropenem, ciprofloxacin, and ceftazidime. Data representative of N≥8 and is presented as mean+/- SD. (Unpaired two-tailed Student’s t-test for column graphs, Two-way ANOVA for grouped data, Mann-Whitney U-test for animal experiment data (*** p < 0.0001, *** p < 0.001, ** p<0.01, * p<0.05))

### Deletion of *rpoE* from STM Δ*pflB* reversed the enhanced antibiotic susceptibility, associated with a recovery in the transcript levels of *acrB* and *tolC*

In *E. coli*, RpoE (σ^E^) functions as an extra cytoplasmic stress sigma factor essential for viability and for coordinating the cellular response to membrane stress (40). The membrane protein RseA, together with the periplasmic factor RseB, binds and sequesters σ^E^ under non-stress conditions, preventing its interaction with RNA polymerase (41, 42). When envelope stress occurs, RseA—which is intrinsically unstable—is rapidly degraded, releasing σ^E^ to activate its target genes. These genes support the synthesis, transport, and assembly of lipopolysaccharides, phospholipids, and outer membrane proteins (OMPs), as well as proteases and chaperones required to preserve or restore outer membrane integrity (42–45). Recent studies further demonstrate that σ^E^ indirectly increases the transcription of the sRNAs *csrB* and *csrC* through σ⁷⁰-dependent promoters (46, 47)

In our previous work, we demonstrated that deletion of *pflB* disrupts intracellular pH homeostasis, leading to membrane depolarization. This perturbation activated the extracytoplasmic sigma factor RpoE, which in turn upregulated transcription of the small RNA *csrB*. The *csrB* RNA can antagonize the activity of the global regulator CsrA by direct binding, thereby impairing processes such as flagellar biosynthesis (10). Notably, CsrA has been shown to stabilize the *acrAB* transcript by interacting with its 5′ untranslated region (48). Given the increased membrane depolarization observed in STM Δ*pflB* upon antibiotic exposure, we hypothesized that RpoE may regulate *acrB* expression indirectly through modulation of the CsrA–*csrB* regulatory circuit.

To test this, we exposed the STM Δ*pflB*Δ*rpoE* strain to sublethal concentrations of meropenem and ciprofloxacin. Compared to STM Δ*pflB*, the double knockout exhibited a significantly reduced proportion of PI-positive cells **(Figure 5A, B)**. Interestingly, deletion of *rpoE* in the STM Δ*pflB* background did not restore membrane potential under basal conditions. However, upon antibiotic challenge, we observed a significant reduction in DiBAC₄-positive cells, indicative of partial recovery in membrane polarization **(Figure 5C, S5A)**. However STM Δ*pflB*Δ*rpoE* had a decreased ROS positive cells compared to STM Δ*pflB*, as assessed using DCF_2_DA staining **(Figure 5D, S5B)**. This was consistent with a decrease in the heightened ipH of STM Δ*pflB* following treatment with ciprofloxacin and meropenem when *rpoE* was simultaneously deleted **(Figure S4B)**.

**Figure 5:**
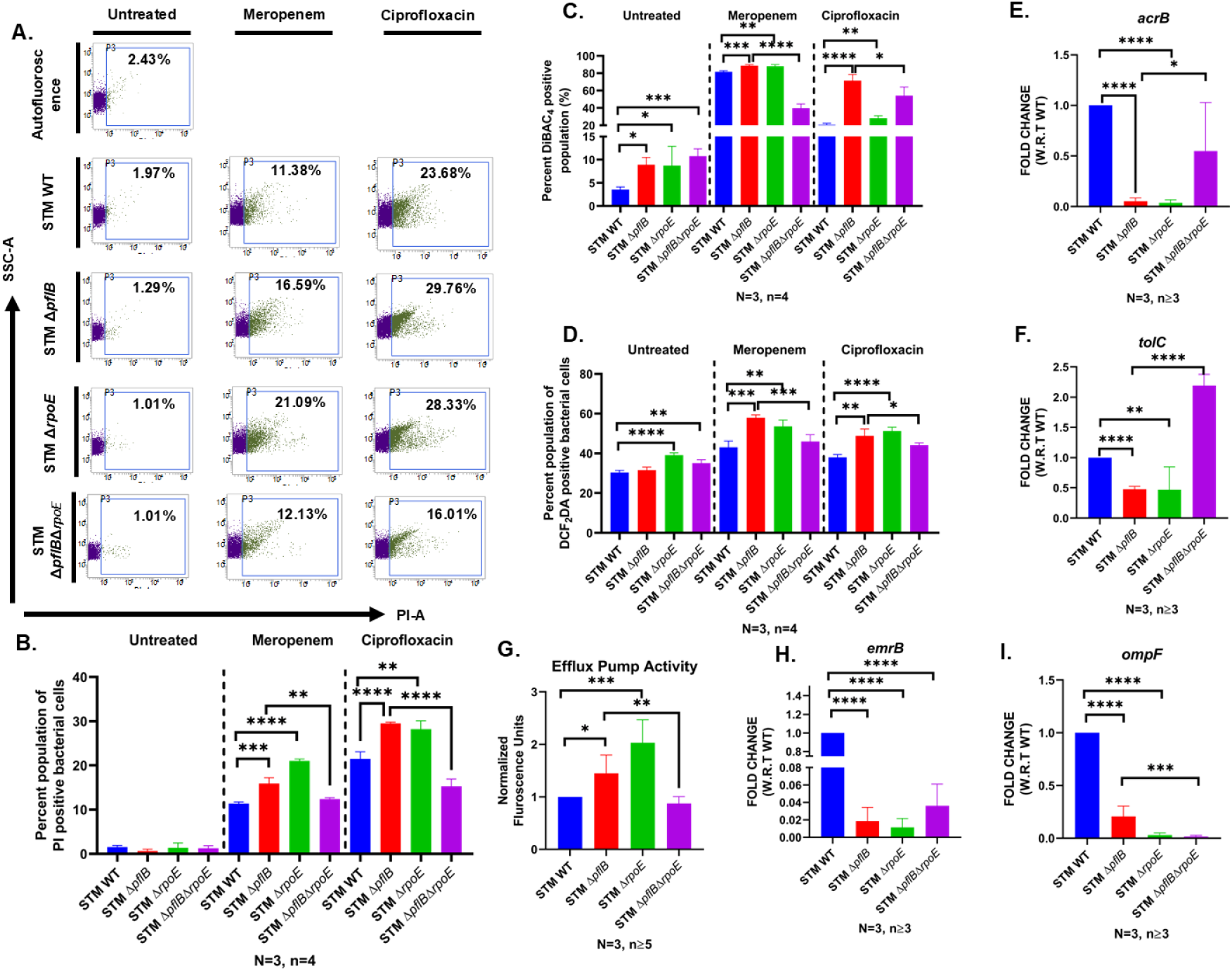
Deletion of *rpoE* in STM Δ*pflB* reversed its increased antibiotic sensitivity, accompanied by restored *acrB* and *tolC* transcript levels. (A) Dot plots (SSC-A vs PI-A) showing antibiotic-mediated killing in STM Δ*pflB* upon *rpoE* deletion. Data representative of N=3, n=4. (B) Bar graphs showing PI positive population in STM Δ*pflB* upon *rpoE* deletion and upon exposure to sub-lethal doses of meropenem and ciprofloxacin. Data representative of N=3, n=4 and is presented as mean+/- SD. (C) Bar graphs display the DiBAC_4_-positive population under meropenem and ciprofloxacin treatment after *rpoE* deletion in STM Δ*pflB*. Data represent N=3, n=4 and are shown as mean ± SD. (D) Bar graphs present the DCF_2_DA-positive population upon exposure to meropenem and ciprofloxacin following *rpoE* deletion in STM Δ*pflB.* Data represent N=3, n=4 and are shown as mean ± SD. (E-F) Transcript levels of key efflux pump genes *acrB* and *tolC* were compared between STM Δ*pflB*Δ*rpoE* and STM Δ*pflB*. Data representative of N=3, n≥3 and is presented as mean+/- SD. (G) Efflux activity assessed using the Nile red assay. Data representative of N=3, n≥5 and is presented as mean+/- SD. (H-I) Transcriptional expression of *emrB* and *ompF* was also evaluated. Data representative of N=3, n≥3 and is presented as mean+/- SD. (Unpaired two-tailed Student’s t-test for column graphs, Two-way ANOVA for grouped data, Mann-Whitney U-test for animal experiment data (*** p < 0.0001, *** p < 0.001, ** p<0.01, * p<0.05))

The improved viability of STM Δ*pflB*Δ*rpoE* relative to STM Δ*pflB* under antibiotic stress was further supported by atomic force microscopy, which revealed increased cellular length and height following exposure to both meropenem and ciprofloxacin **(Figure S6A-C)**.

These phenotypic changes correlated with the restoration of *acrB* and *tolC* transcript levels in STM Δ*pflB*Δ*rpoE*, suggesting enhanced efflux activity **(Figure 5E, F)**. This was further confirmed by increased nile red efflux, indicative of improved pump function **(Figure 5G)**.

Given the role of EmrB in fluoroquinolone resistance, we also examined its expression **(Figure 5H)** (49). As previous studies have not identified a direct regulatory relationship between CsrA and *emrB*, we measured *emrB* transcript levels and observed no significant change in STM Δ*pflB*Δ*rpoE* relative to STM Δ*pflB*. These findings suggest that the observed phenotypic rescue is primarily mediated through the CsrA–*csrB* axis. Deletion of *rpoE* in STM Δ*pflB* led to further downregulation of *ompF*, indicating that while *ompF* suppression does not explain the increased antibiotic sensitivity of STM Δ*pflB*, its further downregulation upon *rpoE* deletion may contribute to the restored resistance observed in STM Δ*pflB*Δ*rpoE* **(Figure 5I)**.

A comparable reduction in CFU for all the strains following 10 mM glutathione antioxidant under sublethal meropenem and ciprofloxacin exposure indicates that the observed differences in bacterial killing are attributable to the distinct levels of ROS generated by each antibiotic **(Figure S7A, B)**(50).

### Deletion of *sirA* abolishes the antibiotic susceptibility of STM Δ*pflB*, along with reduction in ROS levels and membrane depolarization

To further confirm that small RNA *csrB* regulates antibiotic susceptibility, we examined an additional upstream regulator of *csrB*. The BarA/SirA two-component system, which is also known to activate the expression of *csrB* and *csrC* small RNAs, has previously been shown to enhance the expression of HilD, a key regulator of genes within the *Salmonella* Pathogenicity Island-1 (51–53). We deleted *sirA* in the STM Δ*pflB* background and used flow cytometry to quantify propidium iodide-positive cells, revealing a marked reduction in dead cells in the STM Δ*pflB*Δ*sirA* strain compared to STM Δ*pflB* **(Figure 6A, B)**. Interestingly, this decrease in cell death did not correspond with a reduction in membrane depolarization, indicating that the reduced antibiotic sensitivity in STM Δ*pflB*Δ*sirA* was independent of changes in membrane potential **(Figure 6C, S8A)**. A drop in DCF_2_DA-positive cells further supported the observation of diminished antibiotic susceptibility following *sirA* deletion in STM Δ*pflB*. Notably, STM Δ*sirA* alone showed an increase in DCF_2_DA-positive cells after exposure to meropenem and ciprofloxacin **(Figure 6D, S8B)**. This prompted an investigation into efflux pump expression. We observed that *sirA* deletion in STM Δ*pflB* led to elevated expression of *acrB*, *tolC*, *emrB*, and *ompF* **(Figure 6 E-H)**. A nile red assay confirmed restored efflux pump activity in STM Δ*pflB*Δ*sirA* **(Figure 6I)**. Interestingly, deletion of *sirA* alone resulted in decreased *acrB* transcript levels, which may explain the increased DCF_2_DA-positive population upon treatment with meropenem and ciprofloxacin in STM Δ*sirA* **(Figure 6 E)**. However, no overall change in efflux activity was detected in STM Δ*sirA*, possibly due to compensatory roles by other efflux pumps **(Figure 6I)**. These findings suggest that targeting upstream regulators of *csrB* can effectively reverse the antibiotic susceptibility observed in STM Δ*pflB*.

**Figure 6:**
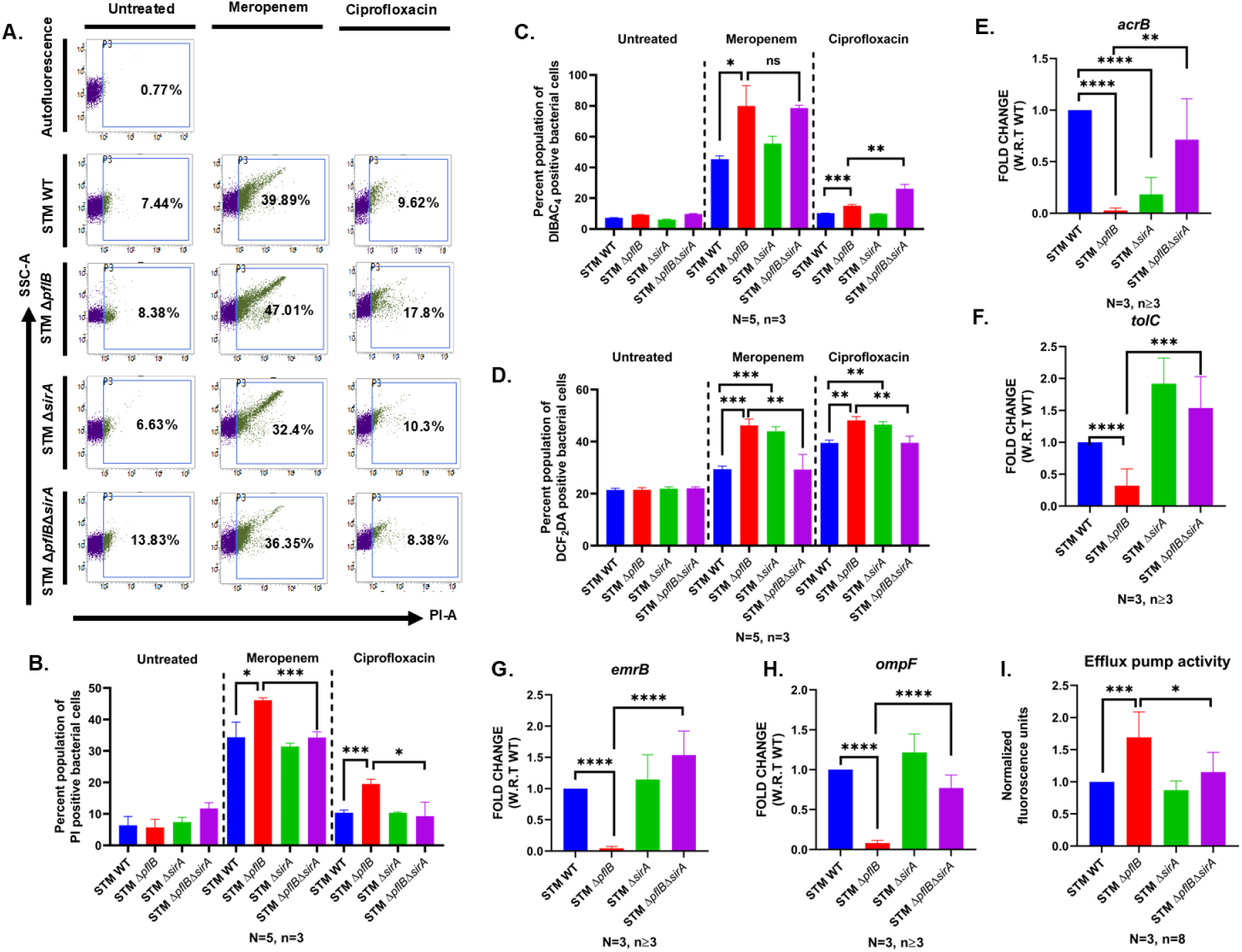
Deleting *sirA* in STM Δ*pflB* similarly reversed its heightened antibiotic sensitivity and restored *acrB* and *tolC* expression levels. (A) Dot plots (SSC-A vs PI-A) showing reduced antibiotic-mediated killing in STM Δ*pflB* upon *sirA* deletion. Data representative of N=5, n=3. (B) Bar graphs showing antibiotic-mediated killing in STM Δ*pflB* upon *sirA* deletion. Data representative of N=5, n=3 and is presented as mean+/- SD. (C) Bar graphs showing DiBAC_4_ population in STM Δ*pflB* upon *sirA* deletion and exposure to antibiotics. Data representative of N=5, n=3 and is presented as mean+/- SD. (D) Bar graphs showing DCF_2_DA population in STM Δ*pflB* upon *sirA* deletion and exposure to antibiotics. Data representative of N=5, n=3 and is presented as mean+/- SD. (E-H) Transcript levels of efflux pump genes *acrB, tolC*, *emrB,* and *ompF* were compared between STM Δ*pflB*Δ*sirA* and STM Δ*pflB*. Data representative of N=3, n≥3 and is presented as mean+/- SD. (I) Efflux activity assessed using the Nile red assay. Data representative of N=3, n=8 and is presented as mean+/- SD. (Unpaired two-tailed Student’s t-test for column graphs, Two-way ANOVA for grouped data, Mann-Whitney U-test for animal experiment data (*** p < 0.0001, *** p < 0.001, ** p<0.01, * p<0.05))

### Deletion of *sirA* ameliorates the heightened susceptibility of STM WT to antibiotics upon exogenous F supplementation

The BarA/SirA two-component regulatory system functions by directly promoting increased expression of the small regulatory RNA *csrB* (54). Given that CsrA is a key regulator of *acrAB* transcript expression, we aimed to determine whether formate-induced signalling through the BarA/SirA system contributes to the enhanced antibiotic sensitivity observed in STM WT (48). To explore this, we assessed the antibiotic susceptibility of the STM Δ*sirA* in the presence of formate.

Formate exposure in STM WT resulted in reduced transcript levels of other efflux-related genes, such as *tolC* and *emrB*. The absence of *sirA* mitigated the suppression of efflux pump gene expression induced by formate in STM WT. Transcript analysis revealed that formate supplementation improved *acrB* expression in STM Δ*sirA*, contrasting with the downregulation of *acrB* seen in formate-treated STM WT **(Figure S9 A-C)**. This reflected in efflux pump activity as well **(Figure S9 D)**.

Consistent with previous data, STM WT treated with formate showed a higher percentage of PI-positive cells following exposure to meropenem and ciprofloxacin. Although STM Δ*sirA* showed a PI-positive population comparable to STM WT when exposed to these antibiotics, adding formate did not further increase PI-positive cells in the mutant under ciprofloxacin treatment. In the case of meropenem treatment, a decrease in extent of PI-positive population was observed **(Figure S9 E**). Likewise, formate treatment did not elevate the ROS-positive population in STM Δ*sirA*, unlike the increase observed in STM WT **(Figure S9 F)**. A similar trend was seen in the number of depolarized cells under antibiotic stress **(Figure S9 G)**.

In summary, these results suggest that extracellular formate enhances STM WT susceptibility to meropenem and ciprofloxacin primarily through the BarA/SirA signalling system.

### Deletion of *pflB* impaired the long-term survival of STM under high antibiotic concentrations, a similar effect observed when formate was supplemented to the wild type

Bacterial persistence in the presence of antibiotics, distinct from antibiotic resistance, remains a persistent challenge in effectively eliminating pathogens in clinical settings(55). Since our data demonstrated that *pflB* deletion increased the antibiotic susceptibility of STM, we aimed to investigate its impact on long-term bacterial survival under high antibiotic stress. Log-phase bacterial cultures, with or without formate supplementation, were exposed to 15× and 25× MIC concentrations of meropenem and ciprofloxacin. Exposure to 15× MIC of meropenem led to a rapid decline in bacterial countsin planktonic culture, with STM Δ*pflB* exhibiting the fastest reduction, reaching statistical significance at 24 hours post-treatment **(Figure 7A)**. Formate supplementation restored survival levels of STM Δ*pflB* to those of the wild type, while supplementation in STM WT also caused a significant reduction in CFUs. This effect was specific to planktonic cultures treated with meropenem and was not observed in biofilm forms or in ciprofloxacin-treated planktonic and biofilm cultures **(Figure 7B, C, D)**.

**Figure 7:**
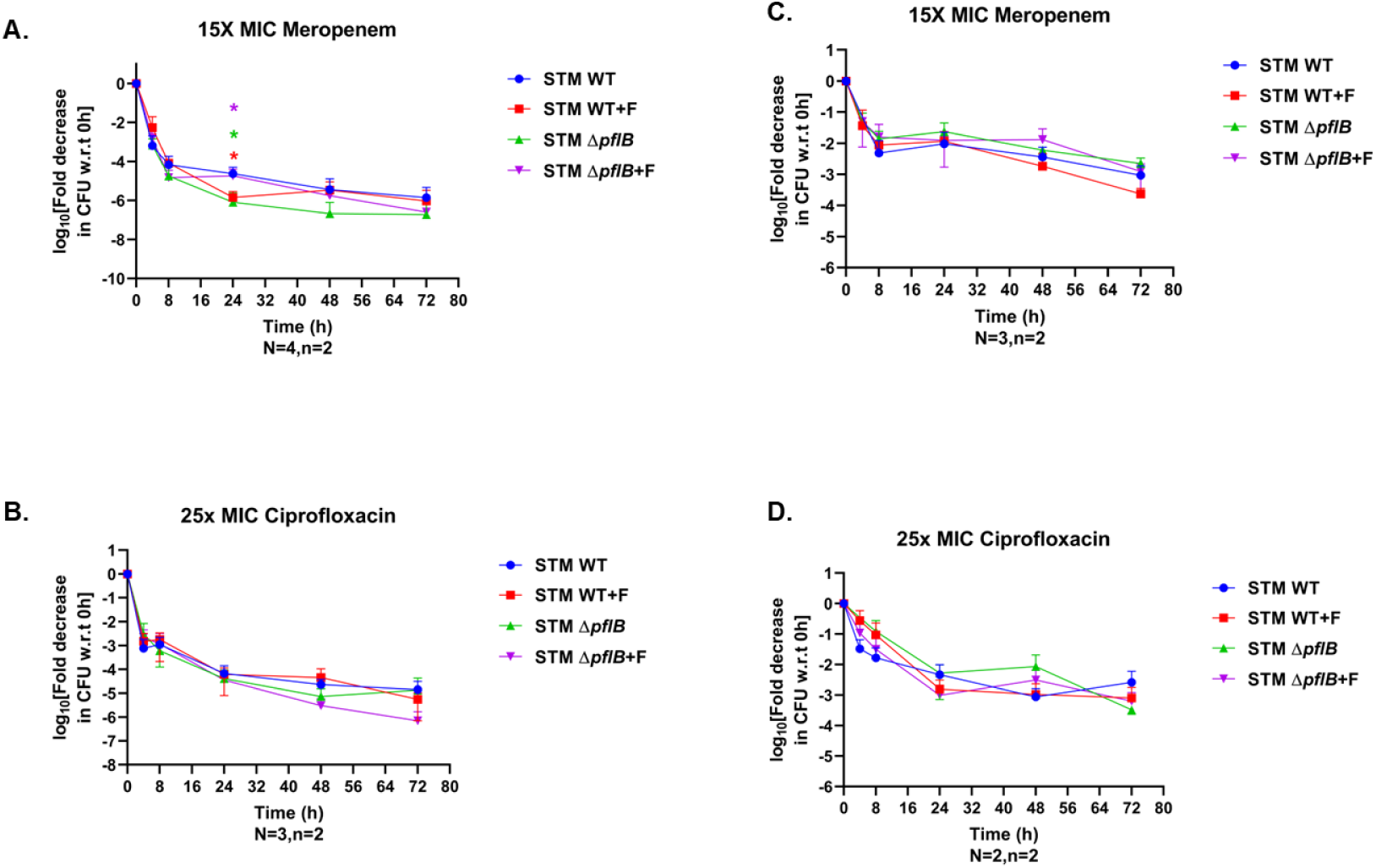
Deletion of *pflB* reduced STM’s long-term survival under high antibiotic concentrations, an effect also seen with formate supplementation in the wild type. (A,B) Line graphs show log₁₀ fold reduction in CFU relative to 0h for planktonic cultures of STM WT and STM Δ*pflB*, with or without formate supplementation, upon exposure to 15X MIC of meropenem and 25X MIC of ciprofloxacin. Data is presented as mean+/- SD. (C, D) Line graphs show log₁₀ fold reduction in CFU relative to 0h for biofilm cultures of STM WT and STM Δ*pflB*, with or without formate supplementation, upon exposure to 15X MIC of meropenem and 25X MIC of ciprofloxacin. Data is presented as mean+/- SD. (Unpaired two-tailed Student’s t-test for column graphs, Two-way ANOVA for grouped data, Mann-Whitney U-test for animal experiment data (*** p < 0.0001, *** p < 0.001, ** p<0.01, * p<0.05))

## Discussion

*Salmonella* has emerged as a major food-borne pathogen with widespread drug resistance, and the increasing prevalence of MDR strains poses a serious clinical challenge (56–58). The growing inefficacy of existing treatments complicates disease management and highlights the urgent need for novel antimicrobial strategies or repurposing of existing drugs (59, 60). Traditionally, research on AMR has focused on canonical molecular mechanisms by which bacteria evade antibiotics, including target modification, enzymatic drug degradation, and altered membrane permeability (61–66).

More recently, bacterial metabolism has gained attention as a determinant of AMR. For example, ceftriaxone-resistant *Salmonella* Derby was shown to exhibit altered metabolic activity, particularly in glutathione biosynthesis, with reduced levels of oxidized glutathione and citrulline. Supplementation of these metabolites restored ceftriaxone susceptibility in MDR isolates (67). Parallel studies have highlighted the influence of ipH on antibiotic resistance in *Salmonella* and other bacteria (11, 15, 16, 68), underscoring the importance of metabolic pathways involved in pH regulation.

The enzyme PflB, which converts pyruvate into formate and acetyl-CoA, plays a central role in maintaining ipH (10). Consistent with this function, we observed that STM lacking *pflB* exhibited increased susceptibility to ciprofloxacin and meropenem, a phenotype that could be partially rescued by external formate supplementation. While pH is known to modulate the PMF and thereby influence efflux activity, our data indicate an additional transcriptional effect, as ipH disruption resulted in downregulation of efflux pump genes *acrB* and *tolC* (11).

In previous work, we showed that membrane stress induced by ipH imbalance activated the RpoE sigma factor, elevating expression of the small RNA *csrB* and repressing flagellar gene expression in STM Δ*pflB* (10). In this study, we extend those findings by showing that *rpoE* deletion not only restored motility but also reversed the heightened antibiotic sensitivity of STM Δ*pflB*. Similarly, deletion of *sirA*, an upstream regulator of *csrB*, had a comparable effect. These results suggest that *pflB* regulates both invasion and antibiotic resistance through a shared RpoE–*csrB* signaling pathway. Importantly, we also found that extracellular formate impacts AMR through the BarA/SirA two-component system—traditionally known for its role in regulating invasion—thereby expanding the functional scope of this signaling axis to include efflux-mediated antibiotic resistance (69) **(Graphical abstract)**.

The role of metabolism in antibiotic tolerance and persistence has also been documented in other bacteria. In *E. coli*, adenosine was shown to suppress the stringent response, enhance PMF, increase respiration, and sensitize persister cells to antibiotics—a process described as “persister awakening” (70). To test whether formate metabolism influences persistence, we assessed long-term survival of STM Δ*pflB* under high-dose antibiotic treatment. In planktonic cultures, STM Δ*pflB* exhibited accelerated killing compared to the wild type, a phenotype that was reversed by formate supplementation. Interestingly, addition of formate to wild-type STM further enhanced antibiotic killing.

Together, our findings support the idea that antibiotic effectiveness is intimately linked to bacterial metabolic state. Specifically, intracellular formate, regulated by PflB, modulates ipH homeostasis, efflux pump expression, and antibiotic tolerance in STM. These results identify formate metabolism as a determinant of both acute resistance and persistence, pointing to metabolic reprogramming as a potential therapeutic strategy to re-sensitize MDR *Salmonella* to existing antibiotics.

## Acknowledgements

We thank the Departmental Real-Time PCR, Divisional Mass Spectrometry, Divisional Flow Cytometry Facility, Divisional Atomic Force Microscopy, and the Central Animal Facility at IISc for supporting experimental work. Dr. Sunitha assisted in formate quantification via GC-MS. Flow cytometry data acquisition was supported by Mr. Munish and Mrs. Navya. Atomic Force Microscopy was supported by Mrs. Monisha. Dr. Raju Rajmani is acknowledged for help in mice serum isolation.

## Author Contributions

DM and DC conceived the study and designed the experiments. DM conducted all experiments, data acquisition, analysis, figure preparation, and wrote the original draft. SVW, PG, and AB helped DM with experiments. DM and DC contributed to manuscript editing.

DC supervised the project and secured funding. All authors reviewed and approved the final manuscript.

## Declaration of Interest

The authors declare no conflicts of interest.

## Funding Information

We gratefully acknowledge financial support from the Department of Biotechnology (DBT) and the Department of Science and Technology (DST), Ministry of Science and Technology. DC thanks DAE for the SRC Outstanding Investigator Award, ASTRA Chair Professorship, and TATA Innovation Fellowship. Additional support came from the DBT-IISc Partnership Program. Infrastructure support from ICMR (Molecular Medicine Center), DST (FIST), and UGC-CAS is also acknowledged. DM is supported by IISc fellowship. PG is supported by DBT-JRF fellowship. AB acknowledges UGC fellowship. The funders had no involvement in study design, data collection, analysis, or publication decisions.

## Data Availability

Data supporting this study are available from the authors upon request.

**Graphical abstract: Graphical abstract showing regulation of AMR in *Salmonella* by formate**

---

**Figure S1:**
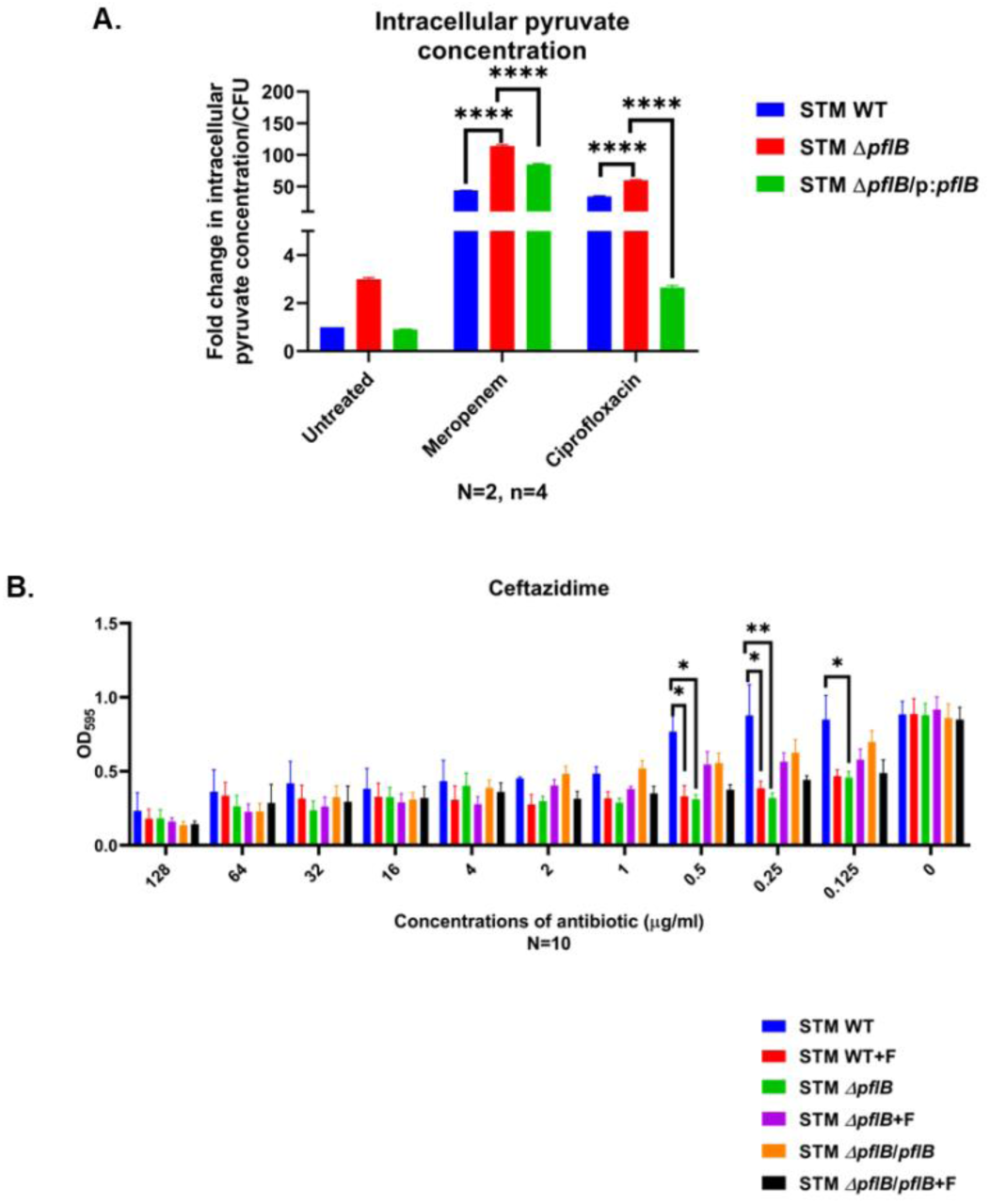
Deletion of *pflB* increased pyruvate levels in STM—a pattern also observed under antibiotic treatment—and additionally heightened STM’s susceptibility to the third-generation cephalosporin ceftazidime, with a comparable sensitivity seen upon extracellular formate supplementation. (A) Bar graphs showing fold change in intracellular pyruvate concentrations in STM WT, STM Δ*pflB*, STM Δ*pflB*/pQE60: *pflB* upon exposure to meropenem and ciprofloxacin. Data representative of N=2, n=4 and is presented as mean+/- SD. (B) Ceftazidime susceptibility in STM WT, STM Δ*pflB*, and STM Δ*pflB*/pQE60: *pflB*, with or without formate, was evaluated using the microfold dilution method. Data is presented as mean+/-SEM of N=10. (Unpaired two-tailed Student’s t-test (column graphs), two-way ANOVA (grouped data), and Mann–Whitney U-test (animal data). (***p < 0.0001, **p < 0.001, **p < 0.01, *p < 0.05))

**Figure S2:**
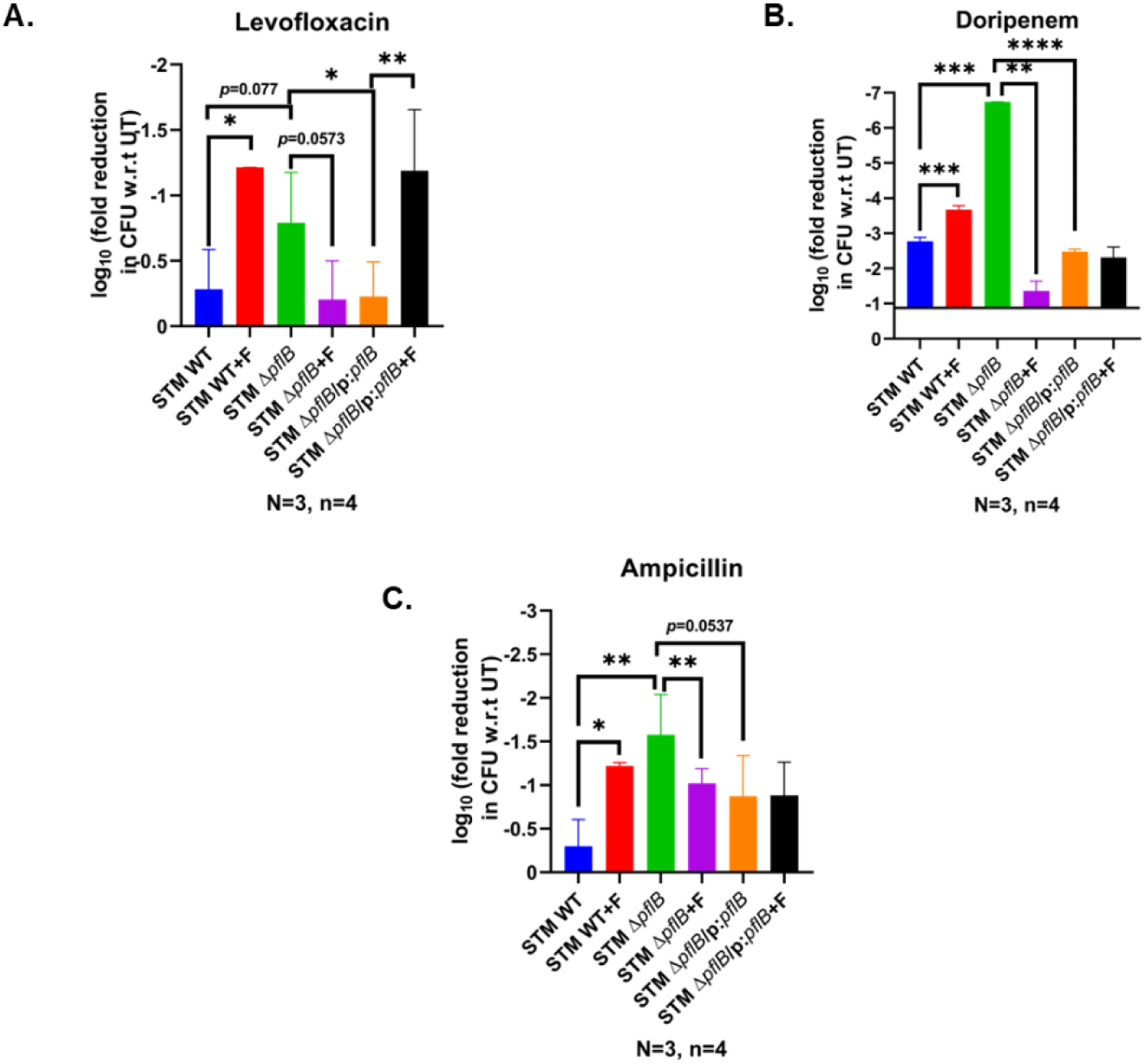
Deletion of *pflB* increased STM susceptibility to the antibiotics levofloxacin, doripenem, and ampicillin, and this effect was partially reversed by formate supplementation. Formate addition to STM WT also heightened its sensitivity to these antibiotics. Bar graphs show the log₁₀ fold reduction in CFU relative to untreated controls for STM WT, STM Δ*pflB*, and STM Δ*pflB*/pQE60: *pflB*, with or without formate, following exposure to sub-lethal concentrations of levofloxacin (A), doripenem (B), and ampicillin (C). Data representative of N=3, n=4 and is presented as mean+/- SD. (Unpaired two-tailed Student’s t-test (column graphs), two-way ANOVA (grouped data), and Mann–Whitney U-test (animal data). (***p < 0.0001, **p < 0.001, **p < 0.01, *p < 0.05))

**Figure S3:**
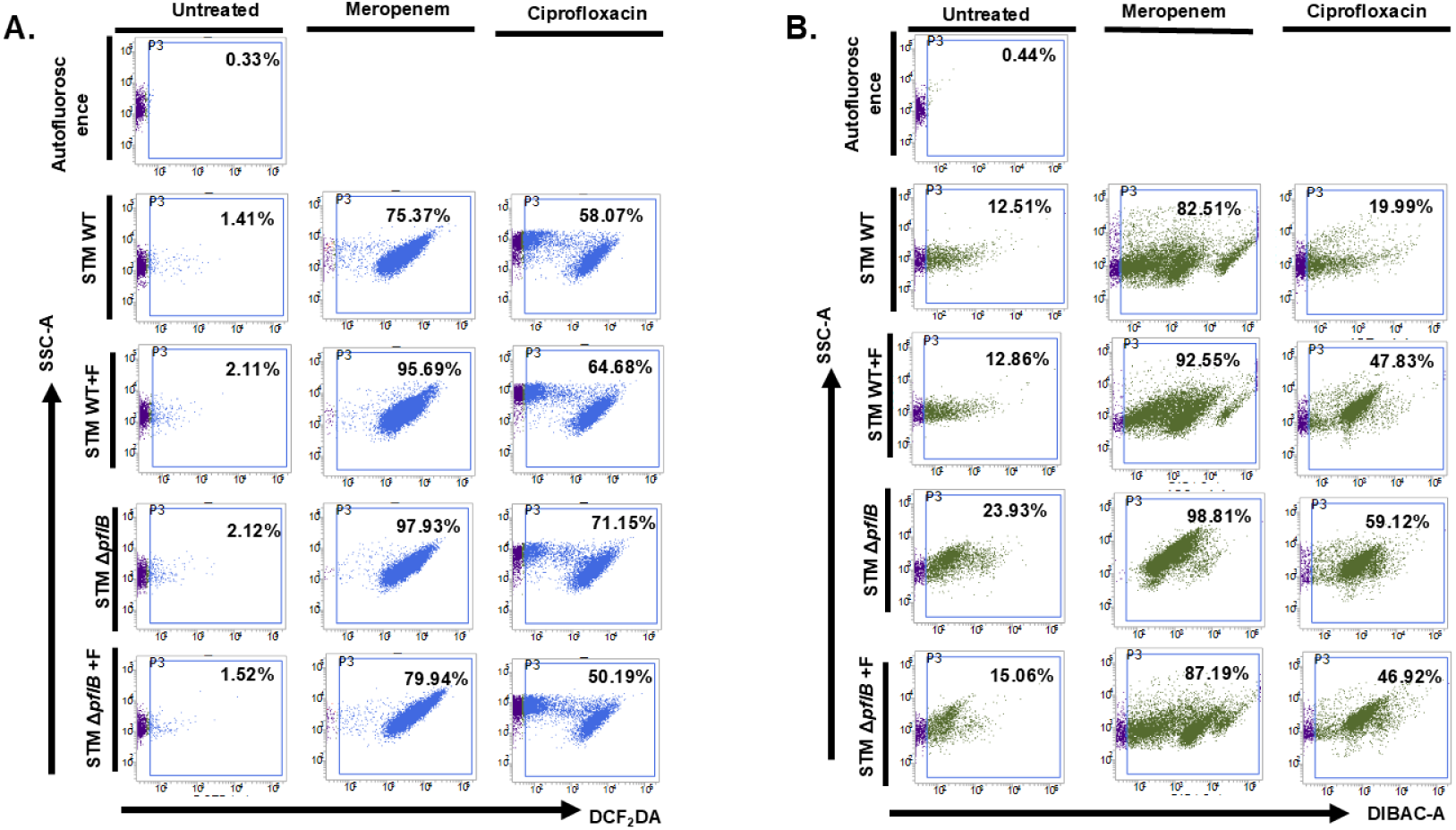
Enhanced antibiotic sensitivity due to *pflB* deletion correlates with increased DCF_2_DA and DiBAC4 staining, indicating elevated ROS levels and membrane depolarization—effects also seen in formate-treated STM WT. (A) Dot plots (SSC-A vs DCF_2_DA) showing intracellular ROS following meropenem and ciprofloxacin treatment in STM WT and STM Δ*pflB* with or without formate supplementation. Data representative of N=3, n≥3. (B) Dot plots (SSC-A vs DiBAC-A) showing membrane depolarization following meropenem and ciprofloxacin treatment in STM WT and STM Δ*pflB* with or without formate supplementation. Data representative of N=3, n≥3. (Unpaired two-tailed Student’s t-test (column graphs), two-way ANOVA (grouped data), and Mann–Whitney U-test (animal data). (***p < 0.0001, **p < 0.001, **p < 0.01, *p < 0.05))

**Figure S4:**
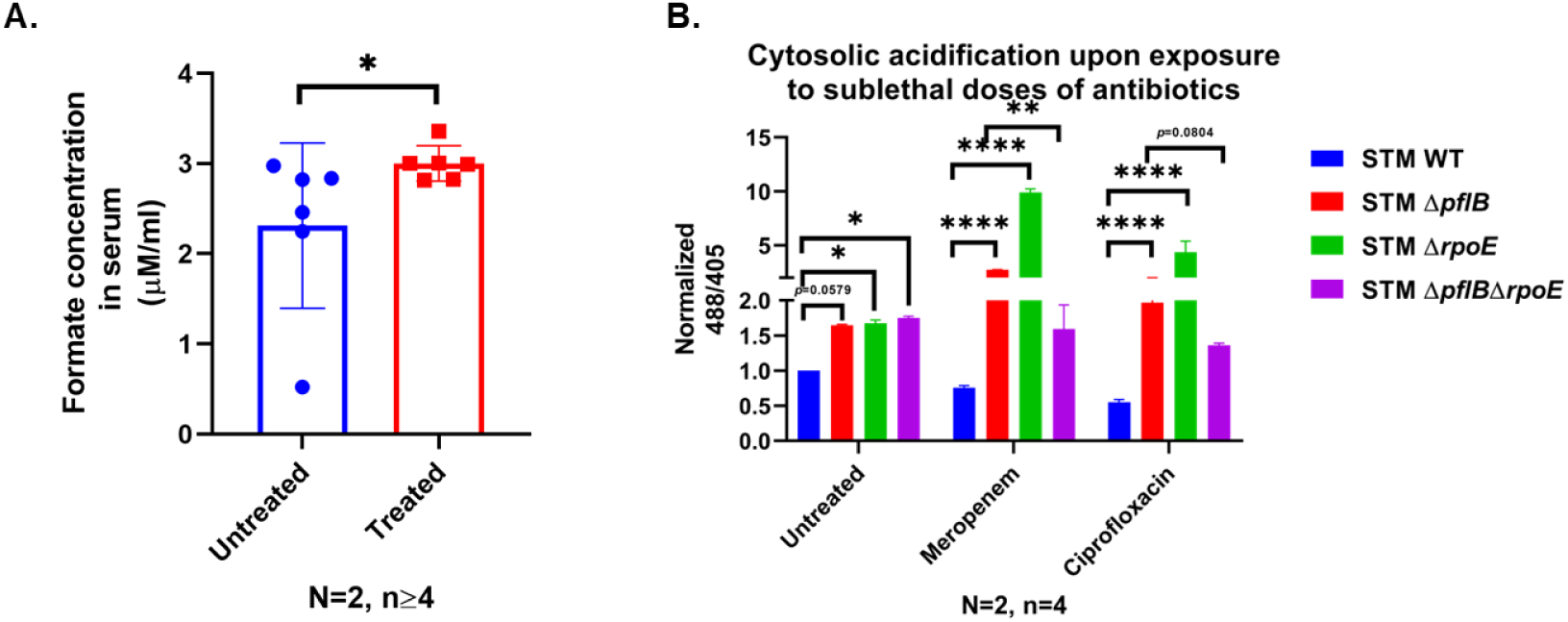
Formate supplementation increased formate levels in serum of treated mice, whereas *rpoE* deletion can also revert the enhanced ipH in STM Δ*pflB* in presence of meropenem and ciprofloxacin. (A) Bar graph showing formate levels in mice serum after exposure to formate through drinking water. (B) Normalized 488/405 ratio of STM WT, STM Δ*pflB*, STM Δ*rpoE*, STM Δ*pflB*Δ*rpoE* in presence of sub-lethal doses of meropenem and ciprofloxacin. Data representative of N=2, n=4 and is presented as mean+/- SD. (Unpaired two-tailed Student’s t-test (column graphs), two-way ANOVA (grouped data), and Mann–Whitney U-test (animal data). (***p < 0.0001, **p < 0.001, **p < 0.01, *p < 0.05))

**Figure S5:**
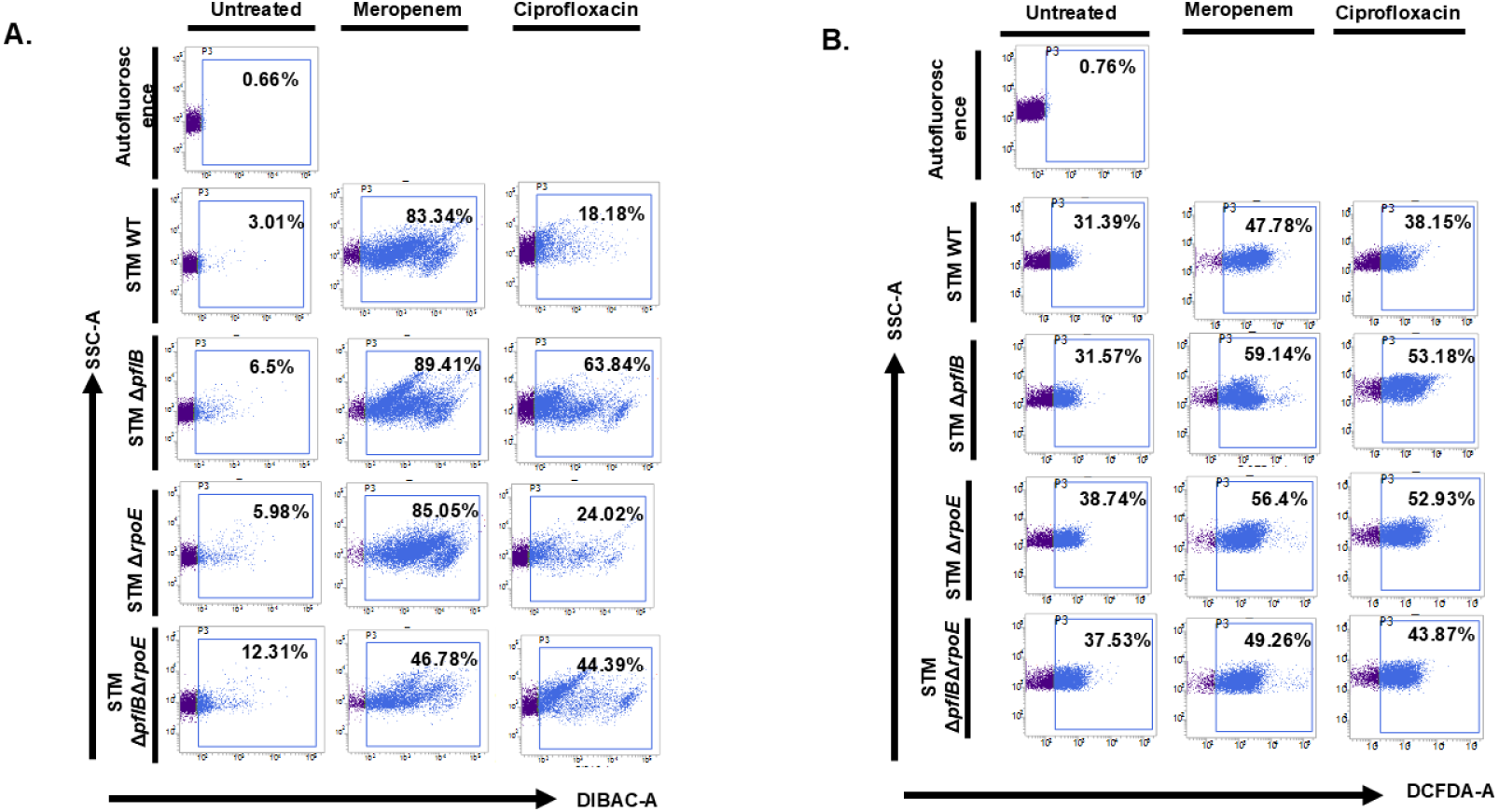
Recovered antibiotic sensitivity due to *rpoE* deletion in STM Δ*pflB* correlates with reduced intracellular ROS levels but it did not have any effect on membrane depolarization. (A) Representative dot plots (SSC-A vs. DIBAC-A) illustrating membrane depolarization in STM WT, STM Δ*pflB*, STM Δ*rpoE,* STM Δ*pflB*Δ*rpoE* strains treated with meropenem or ciprofloxacin. Data shown from three independent experiments (N=3, n=4). (B) Representative dot plots (SSC-A vs. DCFDA-A) showing intracellular ROS in STM WT, STM Δ*pflB*, STM Δ*rpoE,* STM Δ*pflB*Δ*rpoE* following treatment with meropenem or ciprofloxacin. Data representative of N=3, n=4. (Unpaired two-tailed Student’s t-test (column graphs), two-way ANOVA (grouped data), and Mann–Whitney U-test (animal data). (***p < 0.0001, **p < 0.001, **p < 0.01, *p < 0.05))

**Figure S6:**
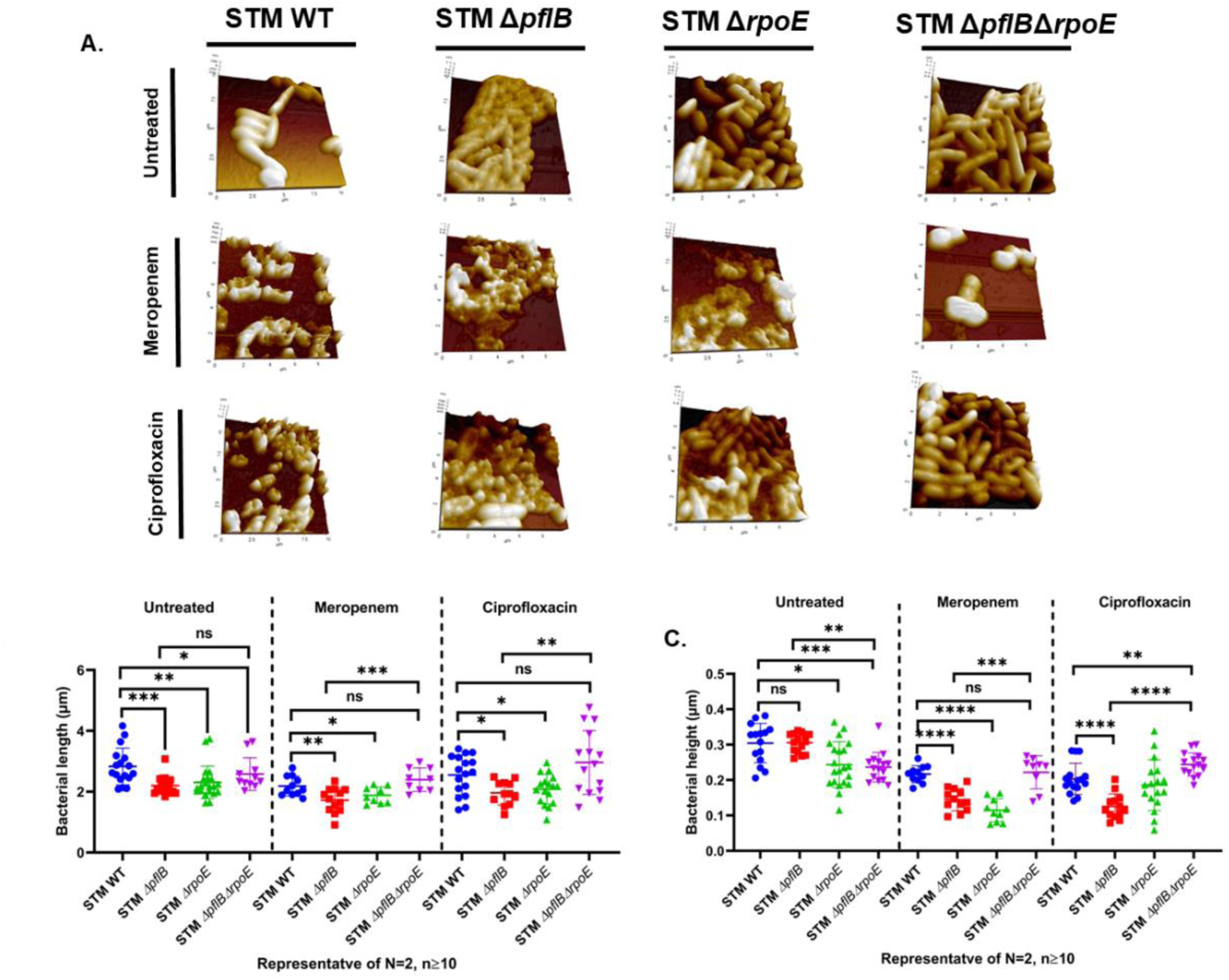
The recovery in antibiotic sensitivity upon *rpoE* deletion in STM Δ*pflB* was also evident from morphological changes under Atomic Force Microscopy. (A) AFM-assissted visualization of morphology of STM WT, STM Δ*pflB*, STM Δ*rpoE*, STM Δ*pflB*Δ*rpoE* upon expsure to sublethal concentrations of meropenem and ciprofloxacin. (B-C) Quantification of bacterial length and height from the same. (Unpaired two-tailed Student’s t-test (column graphs), two-way ANOVA (grouped data), and Mann–Whitney U-test (animal data). (***p < 0.0001, **p < 0.001, **p < 0.01, *p < 0.05))

**Figure S7:**
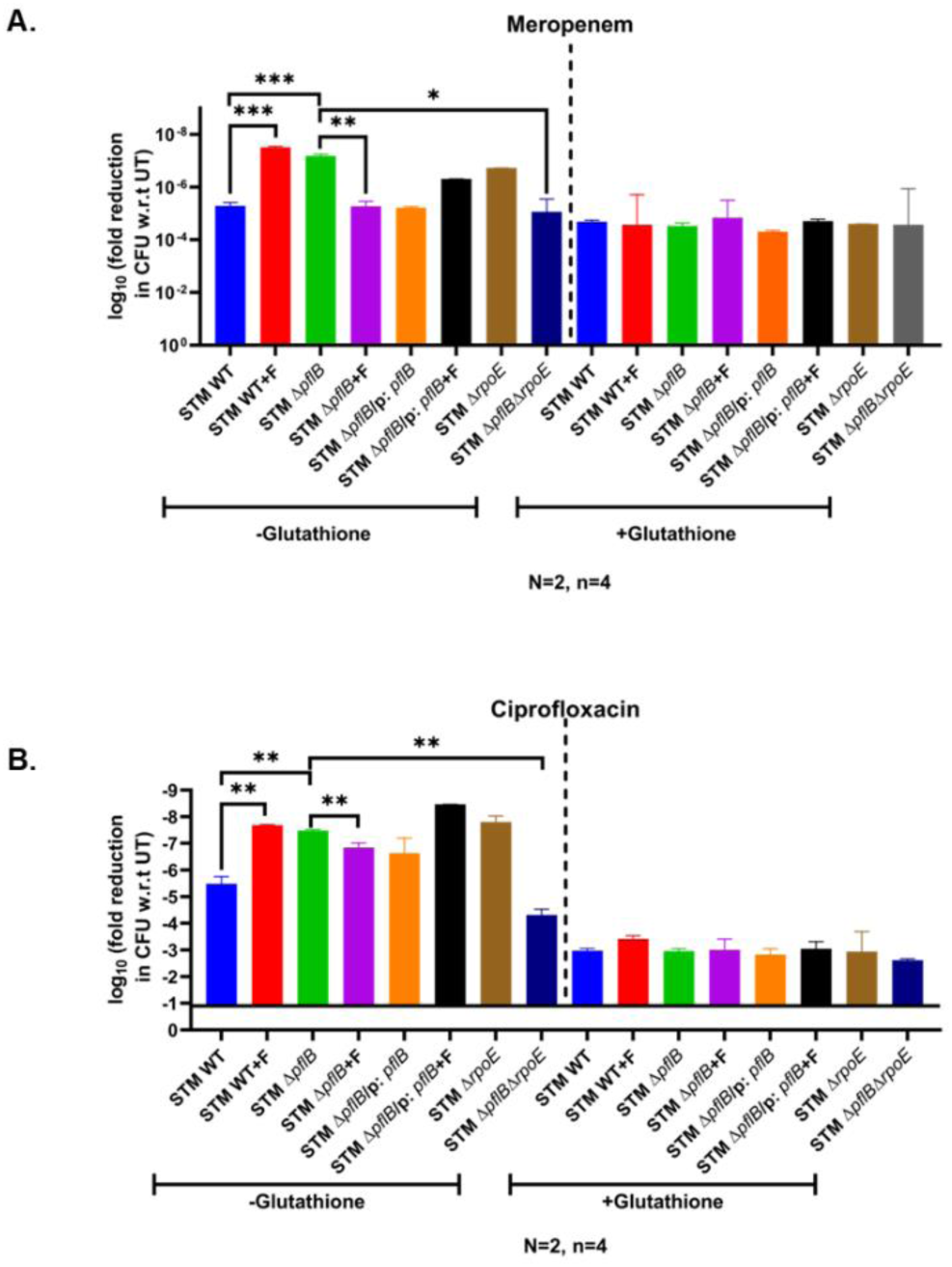
Glutathione treatment equalizes the fold reduction in CFU observed in meropenem- and ciprofloxacin-treated samples. (A-B) Bar graphs showing the log_10_ fold reduction in CFU relative to untreated controls (UT) for STM WT (± formate), STM Δ*pflB* (± formate), STM Δ*pflB*/pQE60: *pflB* (± formate), STM Δ*rpoE*, and STM Δ*pflB*Δ*rpoE* following meropenem or ciprofloxacin treatment, in the presence or absence of glutathione. Data are presented as mean ± SD from two independent experiments (N=2, n=4). (Unpaired two-tailed Student’s t-test (column graphs), two-way ANOVA (grouped data), and Mann–Whitney U-test (animal data). (***p < 0.0001, **p < 0.001, **p < 0.01, *p < 0.05))

**Figure S8:**
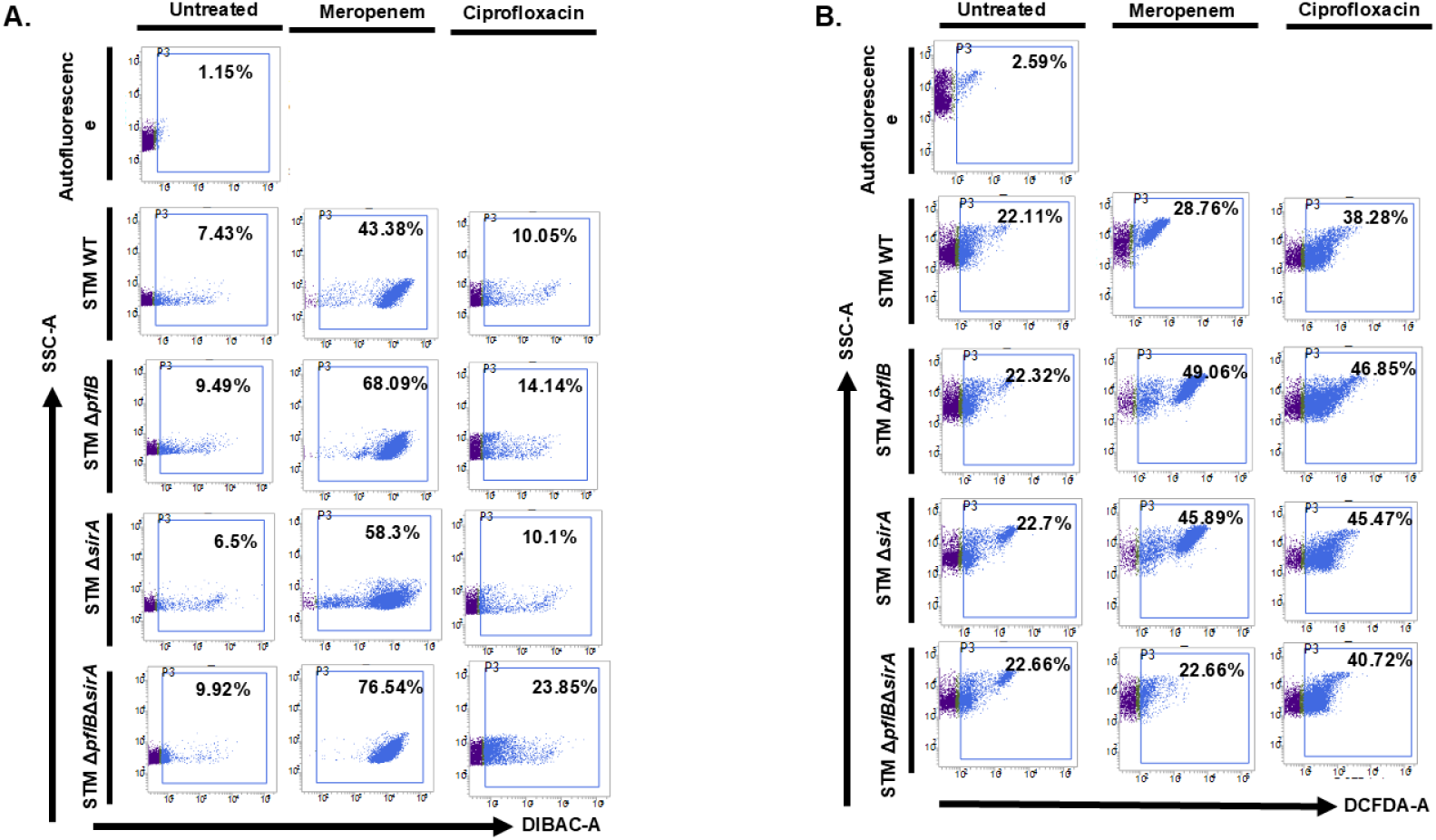
*sirA* deletion mediated reduction of antibiotic susceptibility in STM Δ*pflB* was associated with reduced intracellular ROS levels, but without any effect on membrane depolarization. (A) Representative dot plots (SSC-A vs. DiBAC4) depicting membrane depolarization in STM WT, STM Δ*pflB*, STM Δ*sirA*, and STM Δ*pflB*Δ*sirA* strains following treatment with meropenem or ciprofloxacin. Data shown from five independent experiments (N=5, n=3). (B) Representative dot plots (SSC-A vs. DCFDA-A) showing intracellular ROS levels in STM WT, STM Δ*pflB*, STM Δ*sirA*, and STM Δ*pflB*Δ*sirA* following exposure to meropenem or ciprofloxacin. Data representative of N=5, n=3. (Unpaired two-tailed Student’s t-test (column graphs), two-way ANOVA (grouped data), and Mann–Whitney U-test (animal data). (***p < 0.0001, **p < 0.001, **p < 0.01, *p < 0.05))

**Figure S9:**
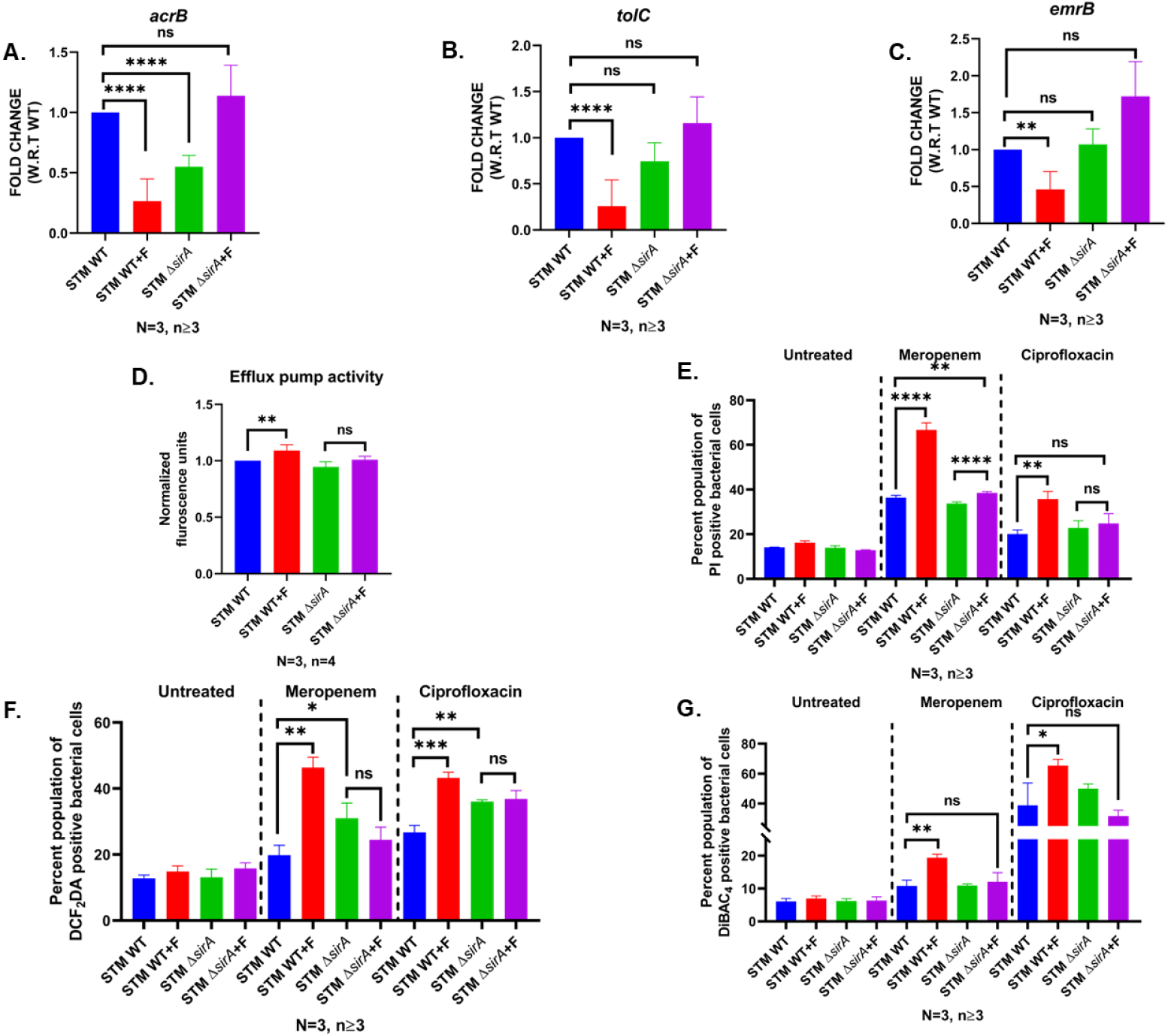
The deletion of *sirA* alleviates the heightened antibiotic sensitivity observed in STM WT upon external formate supplementation. (A-C) RT-qPCR quantification of *acrB*, *tolC*, *emrB*, and *ompF* transcript levels in *STM WT* and STM Δ*sirA*, with or without formate supplementation. Data are presented as mean ± SD from N=3, n≥3. (D) Quantification of PI-positive cells indicating bacterial death under antibiotic stress in STM WT and STM Δ*sirA*, with or without formate supplementation. Data are shown as mean ± SD from N=3, n≥3. (E) Quantification of DCF_2_DA-positive cells in STM WT and STM Δ*sirA* exposed to meropenem or ciprofloxacin, with or without formate supplementation. Representative data of N=3, n≥3 is presented as mean ± SD. (F) Quantification of DiBAC4-positive cells in STM WT and STM Δ*sirA*, with or without formate, reflecting membrane depolarization under antibiotic stress. Representative data of N=3, n≥3 is presented as mean ± SD. (Unpaired two-tailed Student’s t-test (column graphs), two-way ANOVA (grouped data), and Mann–Whitney U-test (animal data). (***p < 0.0001, **p < 0.001, **p < 0.01, *p < 0.05))

## Notes

### Competing Interest Statement

The authors have declared no competing interest.

### Summary of Updates

The manuscript has been updated to include new data.

## References

1. Zhu S, Yang B, Jia Y, Yu F, Wang Z, Liu Y. Comprehensive analysis of disinfectants on the horizontal transfer of antibiotic resistance genes. J Hazard Mater. 2023;453:131428.

2. Brauner A, Fridman O, Gefen O, Balaban NQ. Distinguishing between resistance, tolerance and persistence to antibiotic treatment. Nat Rev Microbiol. 2016;14(5):320–30.

3. Antimicrobial Resistance C. Global burden of bacterial antimicrobial resistance in 2019: a systematic analysis. Lancet. 2022;399(10325):629–55.

4. Azmatullah A, Qamar FN, Thaver D, Zaidi AK, Bhutta ZA. Systematic review of the global epidemiology, clinical and laboratory profile of enteric fever. J Glob Health. 2015;5(2):020407.

5. Kim TH, Hwang HJ, Kim JH. Development of a Novel, Rapid Multiplex Polymerase Chain Reaction Assay for the Detection and Differentiation of Salmonella enterica Serovars Enteritidis and Typhimurium Using Ultra-Fast Convection Polymerase Chain Reaction. Foodborne Pathog Dis. 2017;14(10):580–6.

6. Zhang H, Yao S, Sheng R, Wang J, Li H, Fu Y, et al. A cascade amplification strategy for ultrasensitive Salmonella typhimurium detection based on DNA walker coupling with CRISPR-Cas12a. J Colloid Interface Sci. 2022;625:257–63.

7. Wu S, Ji J, Sheng L, Ye Y, Zhang Y, Sun X. Lysine and valine weaken antibiotic resistance in Salmonella Typhimurium induced by disinfectant stress. J Hazard Mater. 2024;480:135858.

8. Punchihewage-Don AJ, Ranaweera PN, Parveen S. Defense mechanisms of Salmonella against antibiotics: a review. Front Antibiot. 2024;3:1448796.

9. Chowdhury AR, Mukherjee D, Chatterjee R, Chakravortty D. Defying the odds: Determinants of the antimicrobial response of Salmonella Typhi and their interplay. Mol Microbiol. 2024;121(2):213–29.

10. Mukherjee D, Noor S, Mukherjee T, Singh M, Chakravortty D. Intracellular formate modulates a motility-invasion switch in Salmonella Typhimurium. PLoS Pathog. 2025;21(9):e1013453.

11. Lyu Z, Yang A, Villanueva P, Singh A, Ling J. Heterogeneous Flagellar Expression in Single Salmonella Cells Promotes Diversity in Antibiotic Tolerance. mBio. 2021;12(5):e0237421.

12. Chowdhury AR, Mukherjee D, Singh AK, Chakravortty D. Loss of outer membrane protein A (OmpA) impairs the survival of Salmonella Typhimurium by inducing membrane damage in the presence of ceftazidime and meropenem. J Antimicrob Chemother. 2022;77(12):3376–89.

13. Hughes ER, Winter MG, Duerkop BA, Spiga L, Furtado de Carvalho T, Zhu W, et al. Microbial Respiration and Formate Oxidation as Metabolic Signatures of Inflammation-Associated Dysbiosis. Cell Host Microbe. 2017;21(2):208–19.

14. Drescher SPM, Gallo SW, Ferreira PMA, Ferreira CAS, Oliveira SD. Salmonella enterica persister cells form unstable small colony variants after in vitro exposure to ciprofloxacin. Sci Rep. 2019;9(1):7232.

15. Reyes-Fernandez EZ, Schuldiner S. Acidification of Cytoplasm in Escherichia coli Provides a Strategy to Cope with Stress and Facilitates Development of Antibiotic Resistance. Sci Rep. 2020;10(1):9954.

16. Panta PR, Doerrler WT. A link between pH homeostasis and colistin resistance in bacteria. Sci Rep. 2021;11(1):13230.

17. Huang Y, Suyemoto M, Garner CD, Cicconi KM, Altier C. Formate acts as a diffusible signal to induce Salmonella invasion. J Bacteriol. 2008;190(12):4233–41.

18. Papich MG. Saunders handbook of veterinary drugs. North Carolina. 2016;12:162–71.

19. El-Saeed BA, Elshebrawy HA, Zakaria AI, Abdelkhalek A, Sallam KI. Colistin-, cefepime-, and levofloxacin-resistant Salmonella enterica serovars isolated from Egyptian chicken carcasses. Ann Clin Microbiol Antimicrob. 2024;23(1):61.

20. Tang HJ, Chen CC, Zhang CC, Cheng KC, Chiang SR, Chiu YH, et al. Use of Carbapenems against clinical, nontyphoid Salmonella isolates: results from in vitro and in vivo animal studies. Antimicrob Agents Chemother. 2012;56(6):2916–22.

21. Chang HR, Vladoianu IR, Pechere JC. Effects of ampicillin, ceftriaxone, chloramphenicol, pefloxacin and trimethoprim-sulphamethoxazole on Salmonella typhi within human monocyte-derived macrophages. J Antimicrob Chemother. 1990;26(5):689–94.

22. Dwyer DJ, Kohanski MA, Hayete B, Collins JJ. Gyrase inhibitors induce an oxidative damage cellular death pathway in Escherichia coli. Mol Syst Biol. 2007;3:91.

23. Kohanski MA, Dwyer DJ, Hayete B, Lawrence CA, Collins JJ. A common mechanism of cellular death induced by bactericidal antibiotics. Cell. 2007;130(5):797–810.

24. Dwyer DJ, Belenky PA, Yang JH, MacDonald IC, Martell JD, Takahashi N, et al. Antibiotics induce redox-related physiological alterations as part of their lethality. Proc Natl Acad Sci U S A. 2014;111(20):E2100–9.

25. May KL, Grabowicz M. The bacterial outer membrane is an evolving antibiotic barrier. Proc Natl Acad Sci U S A. 2018;115(36):8852–4.

26. Epand RF, Pollard JE, Wright JO, Savage PB, Epand RM. Depolarization, bacterial membrane composition, and the antimicrobial action of ceragenins. Antimicrob Agents Chemother. 2010;54(9):3708–13.

27. Te Winkel JD, Gray DA, Seistrup KH, Hamoen LW, Strahl H. Analysis of Antimicrobial-Triggered Membrane Depolarization Using Voltage Sensitive Dyes. Front Cell Dev Biol. 2016;4:29.

28. Han FF, Gao YH, Luan C, Xie YG, Liu YF, Wang YZ. Comparing bacterial membrane interactions and antimicrobial activity of porcine lactoferricin-derived peptides. J Dairy Sci. 2013;96(6):3471–87.

29. Soren O, Brinch KS, Patel D, Liu Y, Liu A, Coates A, et al. Antimicrobial Peptide Novicidin Synergizes with Rifampin, Ceftriaxone, and Ceftazidime against Antibiotic-Resistant Enterobacteriaceae In Vitro. Antimicrob Agents Chemother. 2015;59(10):6233–40.

30. Cheng M, Huang JX, Ramu S, Butler MS, Cooper MA. Ramoplanin at bactericidal concentrations induces bacterial membrane depolarization in Staphylococcus aureus. Antimicrob Agents Chemother. 2014;58(11):6819–27.

31. Masi M, Refregiers M, Pos KM, Pages JM. Mechanisms of envelope permeability and antibiotic influx and efflux in Gram-negative bacteria. Nat Microbiol. 2017;2:17001.

32. Du D, Wang-Kan X, Neuberger A, van Veen HW, Pos KM, Piddock LJV, et al. Multidrug efflux pumps: structure, function and regulation. Nat Rev Microbiol. 2018;16(9):523–39.

33. Ramaswamy VK, Vargiu AV, Malloci G, Dreier J, Ruggerone P. Molecular Rationale behind the Differential Substrate Specificity of Bacterial RND Multi-Drug Transporters. Sci Rep. 2017;7(1):8075.

34. Rosenberg EY, Ma D, Nikaido H. AcrD of Escherichia coli is an aminoglycoside efflux pump. J Bacteriol. 2000;182(6):1754–6.

35. Webber MA, Bailey AM, Blair JM, Morgan E, Stevens MP, Hinton JC, et al. The global consequence of disruption of the AcrAB-TolC efflux pump in Salmonella enterica includes reduced expression of SPI-1 and other attributes required to infect the host. J Bacteriol. 2009;191(13):4276–85.

36. Ziervogel BK, Roux B. The binding of antibiotics in OmpF porin. Structure. 2013;21(1):76–87.

37. Choi KM, Kim MH, Cai H, Lee YJ, Hong Y, Ryu PY. Salicylic Acid Reduces OmpF Expression, Rendering Salmonella enterica Serovar Typhimurium More Resistant to Cephalosporin Antibiotics. Chonnam Med J. 2018;54(1):17–23.

38. Roy Chowdhury A, Sah S, Varshney U, Chakravortty D. Salmonella Typhimurium outer membrane protein A (OmpA) renders protection from nitrosative stress of macrophages by maintaining the stability of bacterial outer membrane. PLoS Pathog. 2022;18(8):e1010708.

39. Dhayade S, Pietzke M, Wiesheu R, Tait-Mulder J, Athineos D, Sumpton D, et al. Impact of Formate Supplementation on Body Weight and Plasma Amino Acids. Nutrients. 2020;12(8).

40. Hayden JD, Ades SE. The extracytoplasmic stress factor, sigmaE, is required to maintain cell envelope integrity in Escherichia coli. PLoS One. 2008;3(2):e1573.

41. Ahuja N, Korkin D, Chaba R, Cezairliyan BO, Sauer RT, Kim KK, et al. Analyzing the interaction of RseA and RseB, the two negative regulators of the sigmaE envelope stress response, using a combined bioinformatic and experimental strategy. J Biol Chem. 2009;284(8):5403–13.

42. Lima S, Guo MS, Chaba R, Gross CA, Sauer RT. Dual molecular signals mediate the bacterial response to outer-membrane stress. Science. 2013;340(6134):837–41.

43. De Las Penas A, Connolly L, Gross CA. The sigmaE-mediated response to extracytoplasmic stress in Escherichia coli is transduced by RseA and RseB, two negative regulators of sigmaE. Mol Microbiol. 1997;24(2):373–85.

44. Missiakas D, Mayer MP, Lemaire M, Georgopoulos C, Raina S. Modulation of the Escherichia coli sigmaE (RpoE) heat-shock transcription-factor activity by the RseA, RseB and RseC proteins. Mol Microbiol. 1997;24(2):355–71.

45. Campbell EA, Tupy JL, Gruber TM, Wang S, Sharp MM, Gross CA, et al. Crystal structure of Escherichia coli sigmaE with the cytoplasmic domain of its anti-sigma RseA. Mol Cell. 2003;11(4):1067–78.

46. Pannuri A, Vakulskas CA, Zere T, McGibbon LC, Edwards AN, Georgellis D, et al. Circuitry Linking the Catabolite Repression and Csr Global Regulatory Systems of Escherichia coli. J Bacteriol. 2016;198(21):3000–15.

47. Yakhnin H, Aichele R, Ades SE, Romeo T, Babitzke P. Circuitry Linking the Global Csr- and sigma(E)-Dependent Cell Envelope Stress Response Systems. J Bacteriol. 2017;199(23).

48. Ricci V, Attah V, Overton T, Grainger DC, Piddock LJV. CsrA maximizes expression of the AcrAB multidrug resistance transporter. Nucleic Acids Res. 2017;45(22):12798–807.

49. Gu Y, Huang L, Wu C, Huang J, Hao H, Yuan Z, et al. The Evolution of Fluoroquinolone Resistance in Salmonella under Exposure to Sub-Inhibitory Concentration of Enrofloxacin. Int J Mol Sci. 2021;22(22).

50. Parmar K, Kumari Y, Rajmani RS, Chakravortty D. The resilience of *Salmonella* to bile stress is impaired due to the reduced efflux pump activity mediated by the antioxidant enzyme YqhD. bioRxiv. 2024:2024.11.23.625033.

51. Salvail H, Groisman EA. The phosphorelay BarA/SirA activates the non-cognate regulator RcsB in Salmonella enterica. PLoS Genet. 2020;16(5):e1008722.

52. Nava-Galeana J, Nunez C, Bustamante VH. Proteomic analysis reveals the global effect of the BarA/SirA-Csr regulatory cascade in Salmonella Typhimurium grown in conditions that favor the expression of invasion genes. J Proteomics. 2023;286:104960.

53. Gudapaty S, Suzuki K, Wang X, Babitzke P, Romeo T. Regulatory interactions of Csr components: the RNA binding protein CsrA activates csrB transcription in Escherichia coli. J Bacteriol. 2001;183(20):6017–27.

54. Martinez LC, Yakhnin H, Camacho MI, Georgellis D, Babitzke P, Puente JL, et al. Integration of a complex regulatory cascade involving the SirA/BarA and Csr global regulatory systems that controls expression of the Salmonella SPI-1 and SPI-2 virulence regulons through HilD. Mol Microbiol. 2011;80(6):1637–56.

55. Huemer M, Mairpady Shambat S, Brugger SD, Zinkernagel AS. Antibiotic resistance and persistence-Implications for human health and treatment perspectives. EMBO Rep. 2020;21(12):e51034.

56. Algarni S, Ricke SC, Foley SL, Han J. The Dynamics of the Antimicrobial Resistance Mobilome of Salmonella enterica and Related Enteric Bacteria. Front Microbiol. 2022;13:859854.

57. Dyson ZA, Klemm EJ, Palmer S, Dougan G. Antibiotic Resistance and Typhoid. Clin Infect Dis. 2019;68(Suppl 2):S165–S70.

58. Wu S, Yang Y, Wang T, Sun J, Zhang Y, Ji J, et al. Effects of acid, alkaline, cold, and heat environmental stresses on the antibiotic resistance of the Salmonella enterica serovar Typhimurium. Food Research International. 2021;144:110359.

59. Kuang X, Hao H, Dai M, Wang Y, Ahmad I, Liu Z, et al. Serotypes and antimicrobial susceptibility of Salmonella spp. isolated from farm animals in China. Front Microbiol. 2015;6:602.

60. Kebede A, Kemal J, Alemayehu H, Habte Mariam S. Isolation, Identification, and Antibiotic Susceptibility Testing of Salmonella from Slaughtered Bovines and Ovines in Addis Ababa Abattoir Enterprise, Ethiopia: A Cross-Sectional Study. Int J Bacteriol. 2016;2016:3714785.

61. Christou A, Aguera A, Bayona JM, Cytryn E, Fotopoulos V, Lambropoulou D, et al. The potential implications of reclaimed wastewater reuse for irrigation on the agricultural environment: The knowns and unknowns of the fate of antibiotics and antibiotic resistant bacteria and resistance genes - A review. Water Res. 2017;123:448–67.

62. Sunuwar J, Azad RK. A machine learning framework to predict antibiotic resistance traits and yet unknown genes underlying resistance to specific antibiotics in bacterial strains. Brief Bioinform. 2021;22(6).

63. Devanga Ragupathi NK, Muthuirulandi Sethuvel DP, Shankar BA, Munusamy E, Anandan S, Veeraraghavan B. Draft genome sequence of bla(TEM-1)-mediated cephalosporin-resistant Salmonella enterica serovar Typhi from bloodstream infection. J Glob Antimicrob Resist. 2016;7:11–2.

64. Greninger AL, Chatterjee SS, Chan LC, Hamilton SM, Chambers HF, Chiu CY. Whole-Genome Sequencing of Methicillin-Resistant Staphylococcus aureus Resistant to Fifth-Generation Cephalosporins Reveals Potential Non-mecA Mechanisms of Resistance. PLoS One. 2016;11(2):e0149541.

65. Palzkill T. Structural and Mechanistic Basis for Extended-Spectrum Drug-Resistance Mutations in Altering the Specificity of TEM, CTX-M, and KPC beta-lactamases. Front Mol Biosci. 2018;5:16.

66. 66. Sanders CC. beta-Lactamase stability and in vitro activity of oral cephalosporins against strains possessing well-characterized mechanisms of resistance. Antimicrob Agents Chemother. 1989;33(8):1313–7.

67. Ji J, Wu S, Sheng L, Sun J, Ye Y, Zhang Y, et al. Metabolic reprogramming of the glutathione biosynthesis modulates the resistance of Salmonella Derby to ceftriaxone. iScience. 2023;26(8):107263.

68. Wu S, Ji J, Carole NVD, Yang J, Yang Y, Sun J, et al. Combined metabolomics and transcriptomics analysis reveals the mechanism of antibiotic resistance of Salmonella enterica serovar Typhimurium after acidic stress. Food Microbiol. 2023;115:104328.

69. Teplitski M, Goodier RI, Ahmer BM. Pathways leading from BarA/SirA to motility and virulence gene expression in Salmonella. J Bacteriol. 2003;185(24):7257–65.

70. Kitzenberg DA, Lee JS, Mills KB, Kim JS, Liu L, Vazquez-Torres A, et al. Adenosine Awakens Metabolism to Enhance Growth-Independent Killing of Tolerant and Persister Bacteria across Multiple Classes of Antibiotics. mBio. 2022;13(3):e0048022.

